# Rainbow Nucleus Charts Dynamic Interactome of Membrane-less Organelles

**DOI:** 10.1101/2025.05.14.654140

**Authors:** Songtao Ye, Nadav Benhamou Goldfajn, Choon Leng So, Takanari Inoue, Danfeng Cai

## Abstract

Membrane-less organelles (MLOs) perform diverse cellular functions, but how they interact to coordinate these processes remains poorly understood. Here we constructed a “Rainbow Nucleus” cell line to simultaneously visualize five nuclear MLOs using multi-spectral live-cell imaging. We find that MLO interactions are not random: nuclear speckles serve as hubs, and functionally related MLOs interact more frequently than unrelated MLOs. While some interactions are stable, such as those between the histone locus bodies and Cajal bodies, others are transient, such as those between PML nuclear bodies and Cajal bodies. Single particle tracking revealed that stable contacts between Cajal bodies and nuclear speckles are functional, promoting the outward trafficking of small nuclear ribonucleoproteins from Cajal bodies. Finally, RNA Polymerase II-mediated transcription is required to organize the MLO interactome. Our study reveals a structured, transcription-dependent contact network among nuclear MLOs, laying the foundation for future cellular engineering efforts to modulate the MLO interactome.

## Introduction

Membrane-less organelles (MLOs) are dynamic cellular compartments that lack surrounding lipid membranes, yet they play critical roles in organizing biochemical processes within cells^1^. Many MLOs form through liquid-liquid phase separation (LLPS), a biophysical process in which proteins and nucleic acids condense into distinct, liquid-like droplets, enabling localized concentration of reactants and regulation of complex cellular functions^2,3^. In the nucleus, MLOs orchestrate key steps of gene expression, RNA processing, and genome maintenance. For instance, nuclear speckles (NSs) are enriched in splicing factors and involved in mRNA processing; nucleoli are sites of ribosomal RNA transcription and ribosome biogenesis; PML nuclear bodies (PML-NBs) are implicated in antiviral responses, genome stability, and transcriptional regulation; Cajal bodies (CBs) facilitate the maturation of small nuclear RNAs (snRNAs) and small nucleolar RNAs (snoRNAs); and histone locus bodies (HLBs) are specialized for histone gene transcription and mRNA processing^1^. Notably, many nuclear pathways require the coordination of multiple MLOs. For example, snRNAs and snoRNAs mature in CBs before deployment to nuclear speckles and nucleoli, respectively. Additionally, human telomerase reverse transcriptase (hTERT) traffics sequentially through nucleoli and CBs *en route* to telomeres^4^.

While substantial progress has been made in elucidating the molecular mechanisms governing individual MLO formation and function^5–7^, how different MLOs interact and coordinate with one another remains poorly understood^8^. Unraveling these interactions is crucial for understanding the integrated organization of nuclear processes and for uncovering the basis of diseases linked to aberrant MLO dynamics, such as dyskeratosis congenita, a telomere maintenance disorder^9^.

In this study, we developed the **“***Rainbow Nucleus***”** cell line, which simultaneously labels five major nuclear MLOs with spectrally distinct fluorescent proteins in the same cell. We further developed a dedicated analysis pipeline for resolving their dynamics and interactions. This approach reveals how nuclear MLOs interact under physiological conditions and in response to transcriptional inhibition, providing insights into how nuclear organization supports coordinated gene expression and genome regulation.

## Results

### Simultaneous multi-spectral live-cell imaging of five MLOs

To construct the “*Rainbow Nucleus*” cell line, we first used CRISPR/Cas9 to knock-in fluorescent protein tags to the loci of major components of CBs (mStayGold-coilin) and NSs (mScarlet3-SRRM2) (**Figs. 1a-c, S1a-b**) in a HeLa cell line. These endogenously tagged proteins localized to their respective MLOs without disrupting the morphology of MLOs (**Figs. S1c-f**). To visualize the PML-NBs and HLBs, we further stably expressed TagBFP-PML-III and Halo-mini-FLASH (labeled with Janelia Fluorophore 646) respectively at low levels (**Figs. 1a-c, S1f-g**). Finally, the nucleoli were labeled transiently with a long Stokes shift (LSS) – mKate2 attached to nucleophosmin 1 (NPM1), a protein specific to the nucleolus expressed at low levels (**Figs. 1a-c, S1f-g**). The “*Rainbow Nucleus*” achieves five-color imaging with only four excitation laser lines, making it compatible with any standard laser-scanning confocal microscope. This is enabled by the large Stokes shift of LSS-mKate2, which is excited by the 488 nm laser yet emits in a spectral window distinct from mStayGold (**Figs. S1h-k**), allowing both fluorophores to share a single excitation source while remaining separable in emission. The five fluorescent channels were readily separated with minimal spectral crosstalk (**Fig. 1d, Movie S1**), allowing us to capture interactions among MLOs.

**Figure 1.**
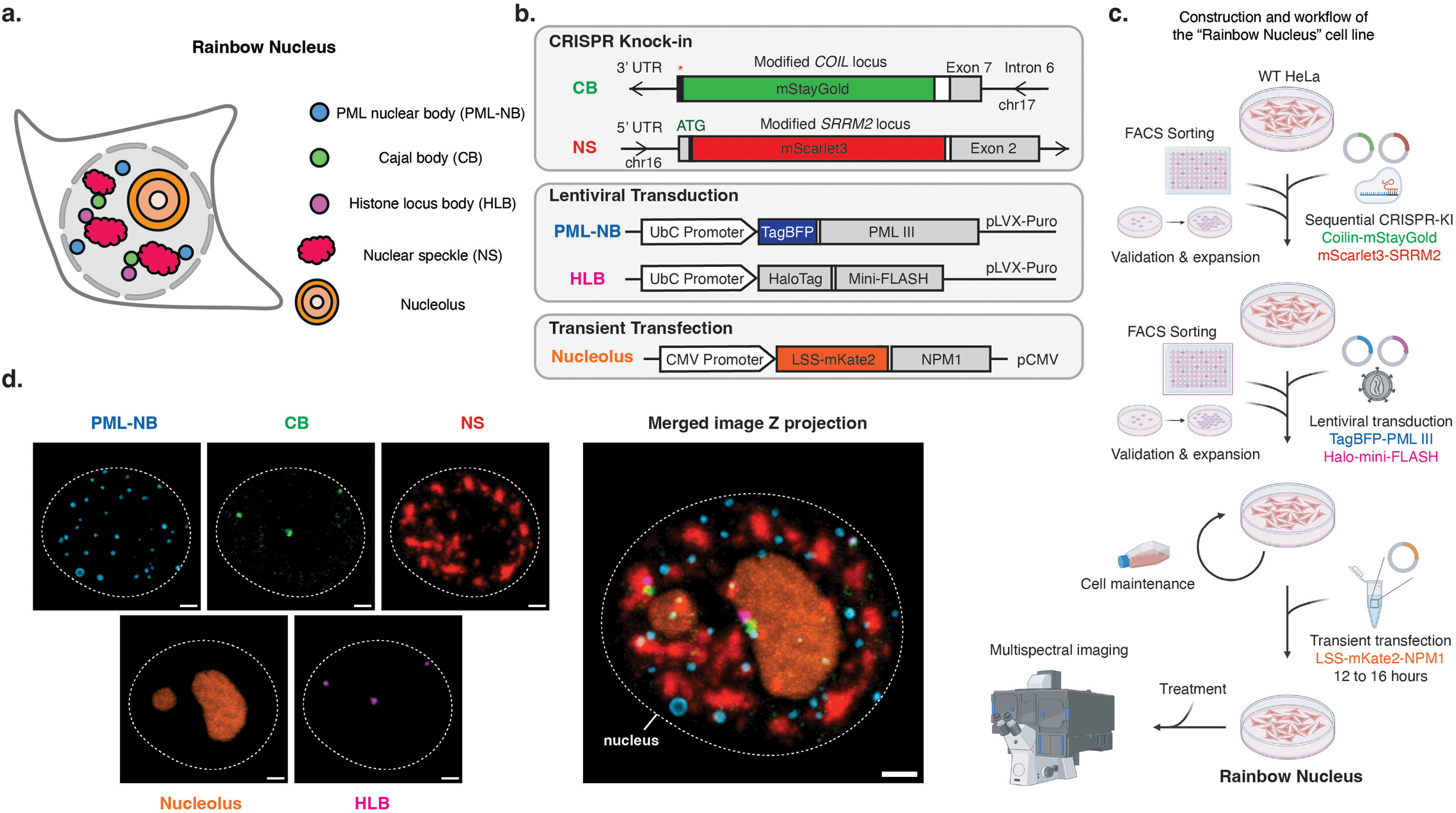
Engineering and validation of the “Rainbow Nucleus” platform for multiplexed imaging of nuclear membrane-less organelles (MLOs). (**a**) Schematic figure of the “Rainbow Nucleus” platform showing the five fluorescently labeled nuclear MLOs: PML nuclear bodies (PML-NBs), Cajal bodies (CBs), histone locus bodies (HLBs), nuclear speckles (NSs), and nucleoli. (**b**) Schematic figure of the genetic approaches for assembling the “Rainbow Nucleus” cell line. Fluorescent tags were introduced through three approaches: CRISPR-mediated knock-in of mStayGold at the *COIL* locus and mScarlet3 at the *SRRM2* locus labeling CB and NS; lentiviral transduction of TagBFP-PML III and HaloTag-mini-FLASH marking PML-NB and HLBs; and transient transfection of LSS-mKate2-NPM1 for nucleolus. (**c**) Schematic figure of the construction workflow of the “Rainbow Nucleus” cell line. The Rainbow Nucleus cell line was constructed through sequential CRISPR knock-in and lentiviral transduction from HeLa cell line, with FACS sorting and validation after each step to ensure correct expression and homogeneity. Cells were maintained at the stage with stable expression of CB, NS, PML-NB and HLB marker, and were transiently transfected with a nucleolus marker prior to imaging. (**d**) Representative maximum intensity projection images of a fixed “Rainbow Nucleus” cell showing individual fluorescent channels for distinct MLOs (left) and the merged image (right). HaloTag protein was labeled with JF 646 ligand. Scale bars, 2 µm.

### 3D imaging of “Rainbow Nucleus” cells reveals nuclear localization and fluidity of all MLOs

We first probed the nuclear distributions of five MLOs using 3D “*Rainbow Nucleus*” (**Fig. 2a**). We found that radially, the nucleoli sat in the center similar to reported before^10^, while other MLOs localized to the periphery, with the PML-NB assuming the outer-most positions (**Figs. 2b, S2a-c**). This is consistent with PML-NBs’ role in organizing heterochromatin domains^11,12^, which are usually close to the nuclear periphery^13–15^. Axially, the distributions of NSs, nucleoli, and HLBs were central and non-polar, which aligns with previous reports of central NS localization^10,16^. In contrast, CBs were shifted upwards, and PML-NBs displayed a significantly broader axial distribution (**Figs. 2b, S2d-f**). Quantification of individual MLO surface area and volume revealed that nucleoli were substantially larger than the other MLOs, whereas NS also reached large sizes but exhibited substantial surface area and volume heterogeneity (**Figs. 2c-d, S2g-j**). Additionally, using live-cell imaging, we saw that the nucleoli moved the slowest, while the PML-NBs moved the fastest among the five MLOs (**Figs. 2e, S2k-l**). The high mobility of PML-NBs is consistent with previous reports describing dynamic behavior of a subset of PML-NBs^17^.

**Figure 2.**
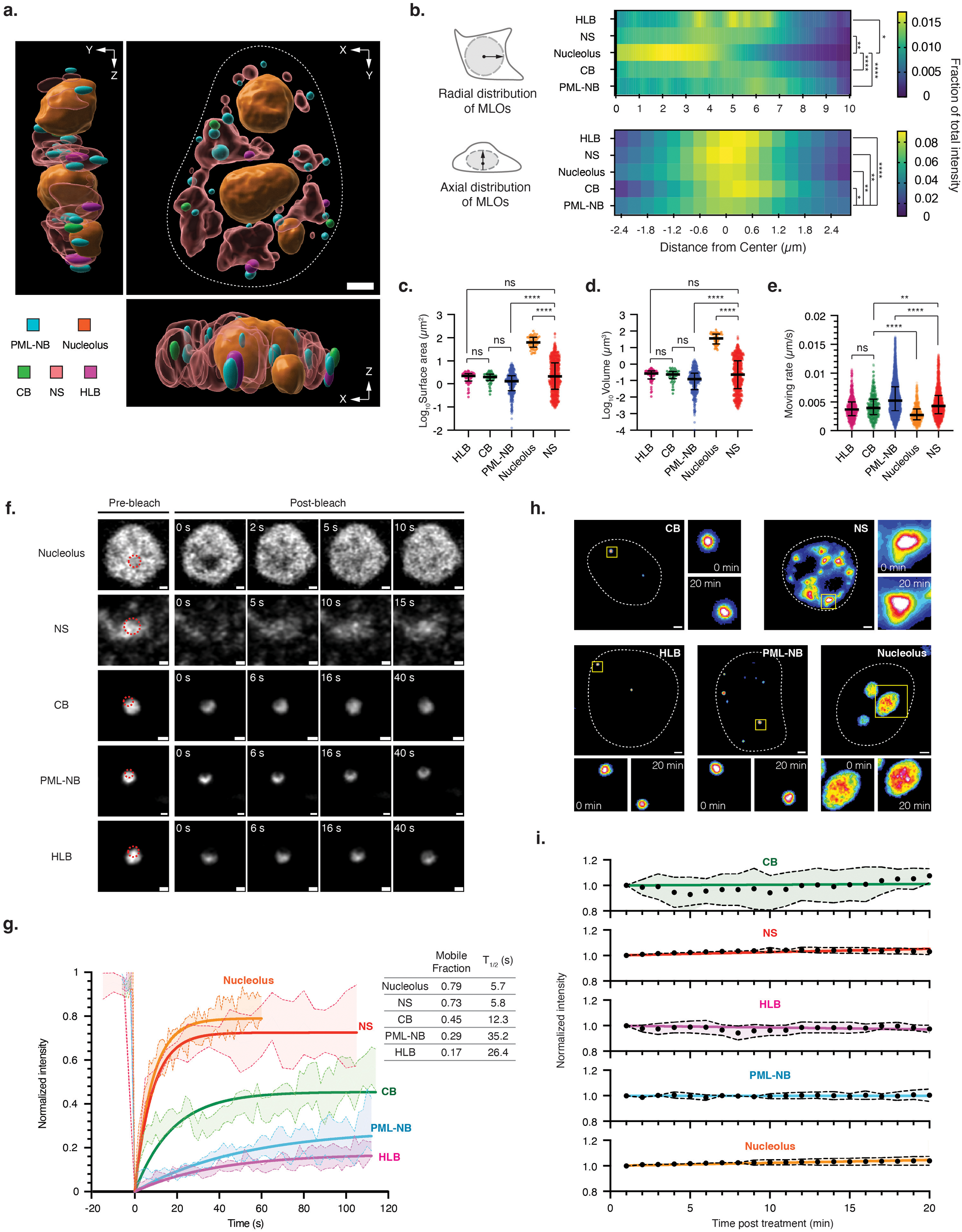
Multispectral imaging reveals spatial organization and dynamics of nuclear MLOs in “Rainbow Nucleus” cells. **(a)** Representative Imaris surface rendering figures showing the spatial organization of nuclear MLOs from a fixed representative “Rainbow Nucleus” cell, visualized along different orthogonal angles (XY, XZ, and YZ). Scale bars, 2 µm. **(b)** Distribution of nuclear MLOs along the radial (top) and axial (bottom) dimensions relative to the center of the nucleus. The nuclear center was defined as the centroid of the NS signals in the radial and axial dimensions. Radial and axial distributions were quantified on a per-cell basis using intensity-over-distance measurements in Imaris software. Statistical comparisons of radial and axial spreads between MLOs were performed using Friedman test followed by Dunn’s multiple comparisons test. Significance levels were annotated as follows: 0.01 < p ≤ 0.05, *; 0.001 < p ≤ 0.01, **; 0.0001 < p ≤ 0.001, ***; p ≤ 0.0001, \*\***\*\***. n = 27 fixed “Rainbow Nucleus” cells. **(c-e)** Scatter plots showing the surface area (**c**), volume (**d**) and moving rate (**e**) of individual nuclear MLO. Horizontal bars indicate medians; vertical whiskers indicate the interquartile range. Data in (**c-d**) represent measurements for individual MLO objects derived from segmentation using Otsu thresholding. n = 52 for HLB, n = 45 for CB, n = 350 for PML-NB, n = 58 for nucleolus, and n = 927 for NS, collected from 27 fixed “Rainbow Nucleus” cells. Moving rates in (**e**) were calculated in Imaris using surface speed function, which calculates instantaneous speed based on changes in MLO object centroid positions across consecutive time points. n = 712 for HLB, n = 494 for CB, n = 8469 for PML-NB, n = 742 for nucleolus, and n = 3206 for NS, collected from 10 live “Rainbow Nucleus” cells across 31 time points (20 s interval; total duration 600 s). Outliers were identified and removed using the ROUT method (Q = 1%). Statistical significance was determined using Kruskal-Wallis tests followed by Dunn’s multiple-comparisons tests. Significance levels were annotated as follows: p > 0.05, ns; 0.001 < p ≤ 0.01, **; p ≤ 0.0001, \*\***\*\***. **(f-g)** Partial fluorescence recovery after photobleaching (partial-FRAP) representative images (**f**) and curves (**g**) of nuclear MLOs in “Rainbow Nucleus” cells. Regions outlined by red dashed circles in (**f**) represent bleaching areas. Scale bars in (**f**), 0.5 µm. Shaded areas and solid lines in (**g**) indicate the standard deviations of the mean values and fitting curves using one-phase association models from FRAP measurements. Inset table in (**g**) summarizes the mobile fraction and half-recovery time (T_1/2_) calculated from curve fitting. All intensities were normalized by min-max scaling, with the pre-bleach fluorescent intensity of the region-of-interest (ROI) set to 1 and the first post-bleach frame set to 0. n = 7 HLBs from 7 live cells, n = 7 CBs from 7 live cells, n = 6 PML-NBs from 6 live cells, n = 10 nucleoli from 10 live cells, and n = 5 NSs from 5 live cells. **(h-i)** Representative images (**h**) and time-dependent fluorescence intensity changes (**i**) of nuclear MLOs upon 1,6-hexanediol treatment in “Rainbow Nucleus” cells. Representative cells are shown in (**h**), with regions outlined by yellow boxes magnified to highlight fluorescence changes at the indicated time points (0 min and 20 min). Scale bars in (**h**), 2 µm. Data points and shaded areas in (**i**) represent the mean normalized intensity values and standard deviations over time, respectively. All fluorescence intensities were normalized to the MLO intensities in the first frame. n = 7 HLBs from 7 live cells, n = 11 CBs from 11 live cells, n = 10 PML-NBs from 10 live cells, n = 11 nucleoli from 11 live cells, and n = 20 NSs from 20 live cells.

Material properties of MLOs are intricately linked to their functions^2,18,19^. “*Rainbow Nucleus*” enables simultaneous measurement of material properties across all five MLOs within the same cell. Using fluorescence recovery after photobleaching (FRAP), we find that the nucleoli and NS are the most dynamic and liquid-like, with T_1/2_ being 5.7 s and 5.8 s, and mobile fractions being 0.79 and 0.73, respectively (**Figs. 2f-g**). On the other hand, CB showed intermediate dynamics (T_1/2_ = 12.3 s; mobile fraction = 0.45), while HLB and PML-NB are the most solid-like with the slowest FRAP recovery (T_1/2_ =26.4 s and 35.2 s; mobile fractions = 0.17 and 0.29) (**Figs. 2f-g)**. However, after 1% 1,6-hexanediol (an aliphatic alcohol that disrupts weak hydrophobic interactions^20^) treatment, the intensity for all MLOs remained constant over 20 min (**Figs. 2h-i**), indicating that complex multivalent forces other than hydrophobic interaction alone govern their formations. Together, these results show that the five nuclear MLOs differ in their material properties, consistent with their distinct assembly mechanisms and cellular functions. While we have established a baseline for MLO material properties, definitively measuring MLO fluidity requires future studies to probe other components using higher-resolution methods like single-particle tracking.

### Global spatial mapping reveals a structured nuclear MLO interactome

To comprehensively map the spatial relationships of MLOs within the nucleus, we developed an integrated analysis pipeline that quantifies all 10 pairwise MLO contact combinations for cumulative probability of encounter (**Figs. 3a, S3a**), median nearest-neighbor surface-to-surface distances (**Fig. 3b**) and contact frequencies (**Fig. 3c**) from a center reference MLO to all other MLOs. These measurements collectively revealed clear, non-random interaction patterns across specific MLO pairs. Previously it has been shown that HLB and CB engage in stable contacts for histone mRNA processing^21^. “*Rainbow nucleus*” confirmed this observation: among all MLO pairs, CB-HLB interactions were the most prominent, reflected by the highest cumulative probability of encountering at shorter distance, a zero median surface-to-surface distance, and high contact frequencies (0.59 and 0.81 when normalized to HLB and CB numbers, respectively), which are significantly higher than randomized pairwise contact expectations (**Figs. 3a-d**). Beyond this strongly enriched CB-HLB pair, distinct and consistent spatial patterns were observed for other MLOs. NS exhibited proximity (**Fig. 3a, S3a, and 3b, first column**) and higher-than-random contact frequencies (**Figs. 3c-d, first column**) to all other MLOs except for the nucleolus. In contrast, the nucleolus is far from other MLOs (**Fig. 3b, second column from left**) and rarely contact with other MLOs, comparable to or below random levels (**Figs. 3c-d, second column from left**). PML-NBs, on the other hand, were close to and contacted frequently with the NS, but separated from and contacted infrequently (near random levels) with CB and HLB (**Figs. 3b-d, middle rows**). Importantly, these pairwise contact patterns were consistent across individual cells (mean cosine similarity = 0.90 ± 0.08; **Fig. S3b**), indicating a highly conserved nuclear MLO interactome across cells. These relationships collectively define a structured nuclear MLO interactome that can be visualized as a network of preferential and depleted interactions (**Fig. 3e**). Within this interactome framework, NS acts as a central interaction hub, having closer proximity and above-random contact frequencies with other MLOs. In contrast, the nucleolus remains comparatively isolated, with larger spatial separations and depleted interactions. The remaining MLOs exhibit selective, pair-specific associations.

**Figure 3.**
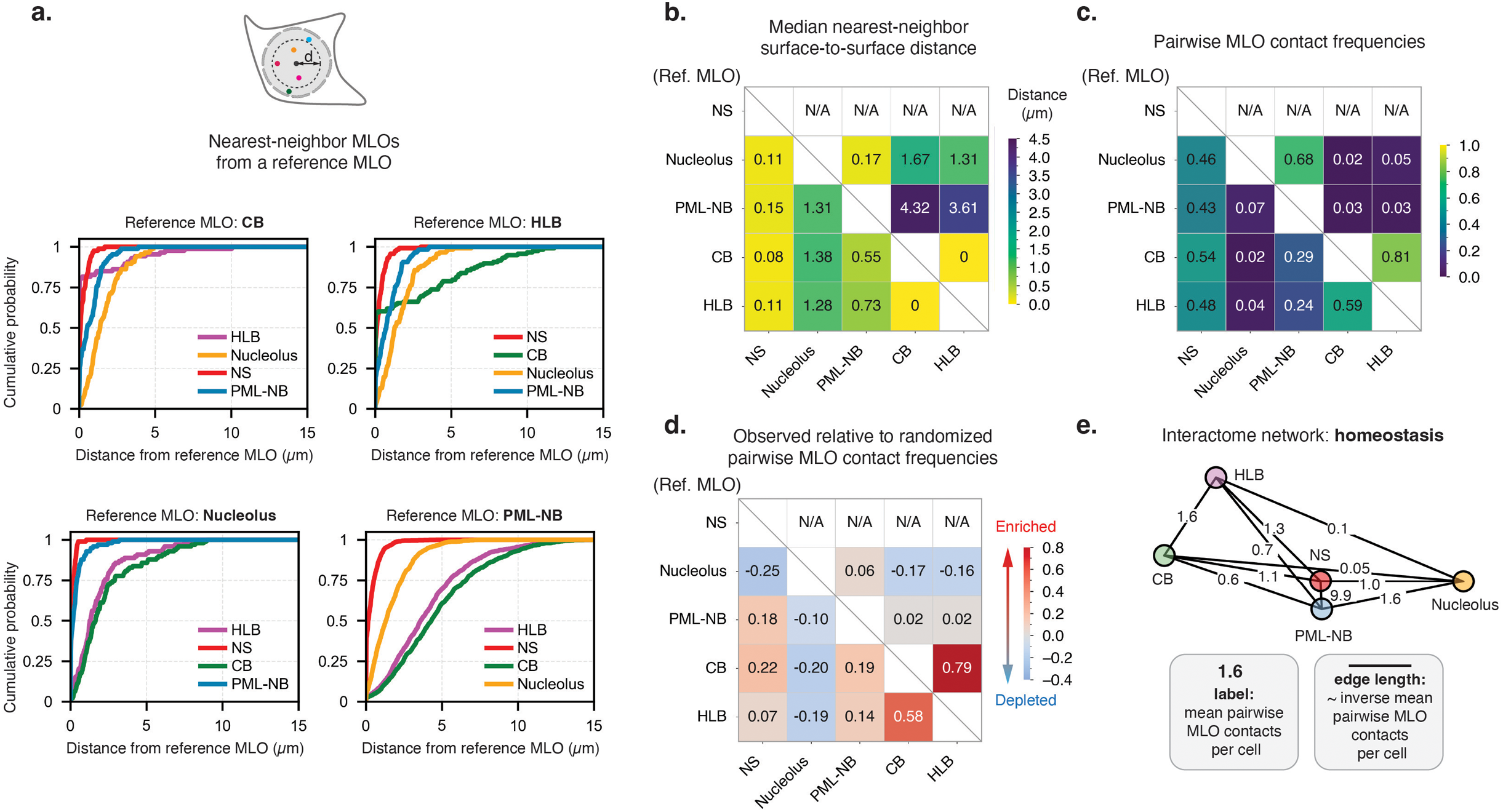
Quantitative mapping reveals a structured, asymmetric interaction landscape among nuclear MLOs. **(a)** Cumulative probability plots indicating the likelihood of reference MLO encountering the nearest MLO of a different type as a function of distance in “Rainbow Nucleus” cells. n = 118 for HLB, n = 87 for CB, n = 987 for PML-NB, and n = 98 for nucleolus, collected from 43 individually 3D-imaged fixed “Rainbow Nucleus” cells. NS was thresholded as a single, whole object per cell in the analysis, as it forms a spatially continuous nuclear compartment rather than discrete punctate structures. Consequently, NS was treated as a target rather than a reference MLO in panel (**a-d**). **(b)** Heatmap of median nearest-neighbor surface-to-surface distances among nuclear MLOs in “Rainbow Nucleus” cells. Color indicates the median distance between each reference MLO (row labels, left) and target MLO (column labels, bottom). n values are the same as in (**a**). **(c)** Heatmap of pairwise MLO contact frequencies among nuclear MLOs in “Rainbow Nucleus” cells. Color indicates the fraction of reference MLOs (row labels, left) in contact with target MLOs (column labels, bottom), relative to the total numbers of reference MLOs. n values are the same as in (**a**). **(d)** Heatmap of observed relative to randomized pairwise MLO contact frequencies among nuclear MLOs in “Rainbow Nucleus” cells. Color indicates the difference between observed and randomized contact frequencies for each pairwise MLO combination, with enrichment (red) and depletion (blue) relative to random expectation. Randomization was performed by repositioning MLO objects within the nuclear volume while preserving their shapes and sizes, and contact frequencies were calculated using the same criteria as in (**c**). n = 10 fixed “Rainbow Nucleus” cells were used for randomization. **(e)** Node-edge network diagram illustrating pairwise interactions among nuclear MLOs under homeostatic conditions in “Rainbow Nucleus” cells. Edge labels indicate the mean counts of pairwise contacts per cell, and edge lengths are inversely proportional to contact frequency. n values are the same as in (**a**).

### Stable and transient interactions exist among different MLOs

While population-wide spatial mapping outlines the architecture of the MLO interactome, it does not distinguish between static proximity and dynamic engagement. To resolve the temporal properties of these behaviors, we tracked individual inter-MLO contacts using multispectral 3D time-lapse imaging (**Figs. 4a-c**), and implemented an integrated analysis pipeline for segmentation, object tracking, and detection of all 10 pairwise MLO contact combinations from these images (**Fig. 4d**). Divergent interaction behaviors emerge, including stable MLO contacts—particularly among HLB-CB, HLB-NS and CB-NS pairs (**Figs. 4a-b, ROI 1**)—as well as transient or no contacts for other MLO pairs (**Figs. 4a-b, ROIs 2-5**). MLO contact kymographs further highlighted the differences in the contact modality (**Fig. 4c**): stable contacts persisted over extended times, often throughout our entire imaging window, with few breakoff events (**Movie S2-3**); whereas transient contacts exhibited frequent, periodic dissociation events on the interaction kymographs, consistent with “kiss-and-run” behavior observed in time-lapse imaging (**Movie S3-4**).

**Figure 4.**
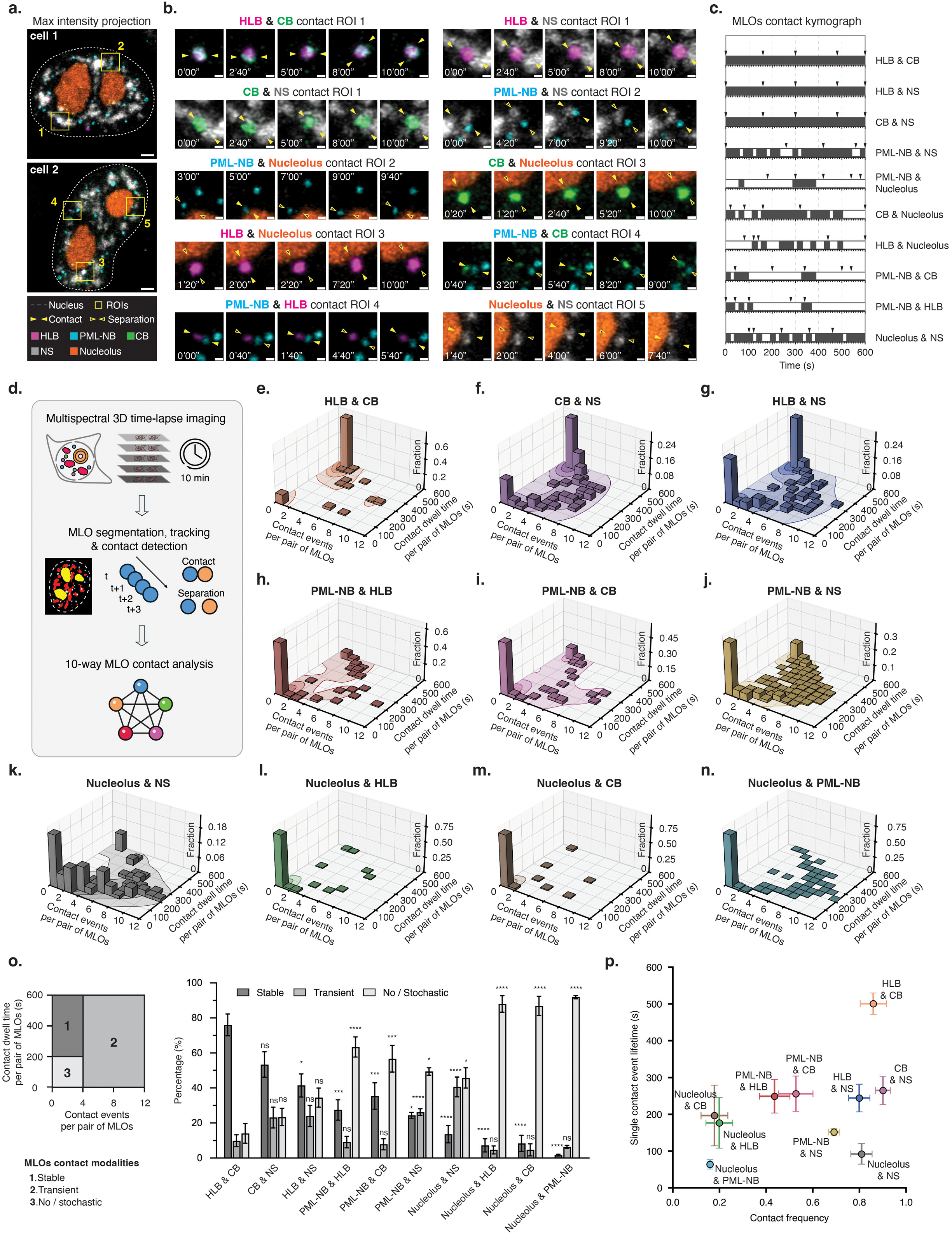
Pairwise nuclear MLO contacts exhibit diverse dynamics that give rise to characteristic distributions of stable and transient contact modalities. **(a-c)** Multispectral 3D time-lapse imaging captures contact states between nuclear MLO pairs in “Rainbow Nucleus” cells. (**a**) Representative maximum intensity projections of two live “Rainbow Nucleus” cells at the initial time point (0 s) of time-lapse imaging. Scale bars, 2 µm. (**b**) Magnified views of the boxed ROIs in (**a**), showing representative contact and separation states across all ten pairwise combinations among five nuclear MLOs. Filled yellow arrowheads indicate contact states, and open arrowheads indicate separation states. Scale bars, 0.5 µm. (**c**) MLO contact kymographs corresponding to the ten pairwise MLO combinations in (**b**). Contact (solid lines) and separation (gaps) states are shown for the indicated MLO pairs. Arrowheads mark the time points selected for display in (**b**). Representative frames in panels (**b-c**) were selected to emphasize clearly distinguishable contact and separation states. **(d)** Schematic figure demonstrating the workflow for determining the contact dynamics of nuclear MLO pairs. Multi-spectral 3D time-lapse imaging is followed by MLO segmentation, tracking, and contact detection to quantify pairwise interaction dynamics across all ten pairwise combinations among five nuclear MLOs. **(e-n)** Probability density plots showing the landscapes of pairwise MLO contact dynamics. Each MLO pair was defined by its contact event counts (X axis) and cumulative contact dwell time (Y axis) during the imaging period. All MLO pairs are collectively represented by density landscapes and column distributions. Contour lines (5%, 25%, 50%, and 75% of maximum density) and filled regions on the XY plane represent relative probability density, and the columns represent the percentage of total contact events within each region. Non-interacting pairs are assigned to (0,0) on the XY plane, with counts calculated by subtracting observed contacts from the total possible contacts (defined as the smaller value of the two MLO populations; for pairs involving NS, the total possible contacts were defined by the population of the non-NS MLO). An exception was made for nucleolus-PML-NB (**n**) pairs, which the PML-NB number was used as total possible contacts due to one single nucleolus can often contact multiple PML-NBs. Sample sizes are as follows: (**e-f, i, m**), n = 65 pairs; (**g-h**), n = 85 pairs; (**j, n**), n = 814 pairs; (**k-l**), n = 82 pairs, collected from 32 live “Rainbow Nucleus” cells imaged over 10 minutes at 20-second intervals. **(o)** Classification and quantification of the pairwise MLO contact modalities. Left panel, schematic figure defining stable, transient and no/stochastic contact modalities based on contact dynamics. Right panel, bar graph showing the percentage of stable, transient, and no/stochastic interaction modes for each pairwise MLO interaction across cells. Data represent mean values with error bars indicate the standard error of mean (SEM). Statistical comparisons for each interaction modality of each pairwise combination were performed against the corresponding modality of HLB & CB using Kruskal-Wallis tests followed by Dunn’s multiple-comparisons tests. Significance levels were annotated as follows: p > 0.05, ns; 0.01 < p ≤ 0.05, *; 0.0001 < p ≤ 0.001, ***; p ≤ 0.0001, \*\***\*\***. Sample sizes are the same as in (**e-n**). **(p)** Scatter plot showing the contact frequency and single contact event lifetime for pairwise MLO interactions. Contact frequency (X axis) is defined as the fraction of observed contacts relative to the total possible contacts per cell for the indicated MLO pair, and single contact event lifetime (Y axis) represents the average duration of uninterrupted contact periods prior to separation. Data represent mean values with error bars indicate SEM. Single contact event lifetime measurements included only cells containing at least one detectable contact event, with n = 32 for CB & NS and PML-NB & NS; n = 31 for HLB & NS and Nucleolus & NS; n = 30 for Nucleolus & PML-NB; n = 29 for HLB & CB; n = 24 for PML-NB & HLB; n = 23 for PML-NB & CB; n = 11 for Nucleolus & HLB; and n = 9 for Nucleolus & CB, collected from 32 live “Rainbow Nucleus” cells imaged over 10 minutes at 20-second intervals. Contact frequency measurements included all of 32 live “Rainbow Nucleus” cells for each MLO pair. Statistical segregation of pairwise MLO contacts was assessed using PERMANOVA based on per-cell contact frequency and single contact event lifetime measurements (pseudo-F = 13.24, p = 0.00001, total 99999 permutations).

Applying the same imaging and analysis pipeline across a larger cell population, we quantified the number of contact events and the total contact dwell times during a 10-min imaging window and aggregated these data in contact kymographs (**Fig. S4**) and contact density maps (**Figs. 4e-n**) to further classify distinct contact modalities. The resulting graphs confirmed the stable contacts among HLB-CB, HLB-NS and CB-NS pairs, clustering in regions with few contact events but long dwell times (**Figs. 4e-g**, top left corner of the density map; **Figs. S4a-c**). In contrast, PML-NB-engaged (**Figs. 4h-j; Figs. S4d-f**) and nucleolus-engaged contacts (**Figs. 4k-n; Figs. S4g-j**) predominantly exhibited higher contact frequencies with a broad distribution of dwell times, indicating they form transient contacts. In particular, the nucleolus contacted little with HLB, CB, or PML-NB, as indicated by minimal overlap in contact occurrence (**Figs. S4h-j, Figs. 4l-n**), suggesting that these nucleolus-engaged contacts are infrequent and short-lived.

We further categorized all these interactions into stable (duration ≥ 200s and ≤ 4 contact events), transient (4-12 contact events), and no/stochastic (duration < 200s and < 4 contact events) interactions, and confirmed that HLB-CB and CB-NS interactions are the most stable, accounting for around 80% and 50% of all interactions, while the nucleolus barely engages in stable interactions with other MLOs (**Fig. 4o**). Consistently, plotting MLO pairs by contact frequency and single-contact lifetime further distinguished their interaction behaviors (**Fig. 4p**). HLB-CB interactions exhibited the longest contact lifetimes, with CB-NS and HLB-NS interactions combined relatively long lifetimes with high contact frequencies. The remaining MLO pairs generally displayed lower contact frequencies and/or shorter contact lifetimes. Both MLO pairwise contact frequencies and single-contact lifetimes were highly conserved across individual cells (mean cosine similarity = 0.97 and 0.72, respectively; **Fig. S5**), indicating that MLO contact dynamics are consistent across cells. In summary, MLOs engage in non-random, specific modes of interactions. HLB, CB, and NS form predominantly stable contacts among themselves, whereas other MLO pairs engage in transient or no/stochastic contacts.

### Stable inter-MLO contacts resist mechanical perturbation

To test if the stable contacts between different MLOs are active connections as opposed to passive ones due to the co-occupancy of the MLOs in the same confined space, we attempted to disrupt the CB-HLB interactions using an ActuAtor tool to apply intracellular force selectively on the nuclear envelope^22^. ActuAtor contains a bioengineered bacterial actin regulator (ActA-FKBP) that dimerizes with FRB-Sec61B when rapamycin is added. This recruits ActA-FKBP to the nuclear outer membrane, where it nucleates actin polymerization, deforming the nucleus and disrupting the intranuclear localization of MLOs (**Fig. 5a**). Consequently, MLOs mediated through active connections will retain their contacts despite location changes, whereas MLO co-occupancy by chance will be disrupted. Upon addition of rapamycin, we observed actin polymerization on the nuclear envelope as tracked by LifeAct signals^22^, accompanied by protrusions and bleb formations on the nucleus, indicative of nuclear deformation (**Fig. 5b, Movie S5**). During nuclear deformation, CB and HLB interacted stably and moved in a highly coordinated manner (**Figs. 5c-d, top panels, Movie S6**). This was reflected by high Pearson correlations for track displacement (**Fig. 5e**) and high cosine similarity for track directions (**Fig. 5f**), suggesting that CBs and HLBs engage in active and stable contacts which can survive mechanical perturbation. In contrast, transient contacts, such as those between PML-NB and CB or HLB, displayed uncoordinated movements and frequent separations (**Figs. 5c-d, bottom panels, Movie S6**), indicated by reduced correlation and directional similarity (**Figs. 5g-h**). These results all indicate that stable interactions such as the CB–HLB association are specific, perturbation-resistant connections rather than incidental colocalization.

**Figure 5.**
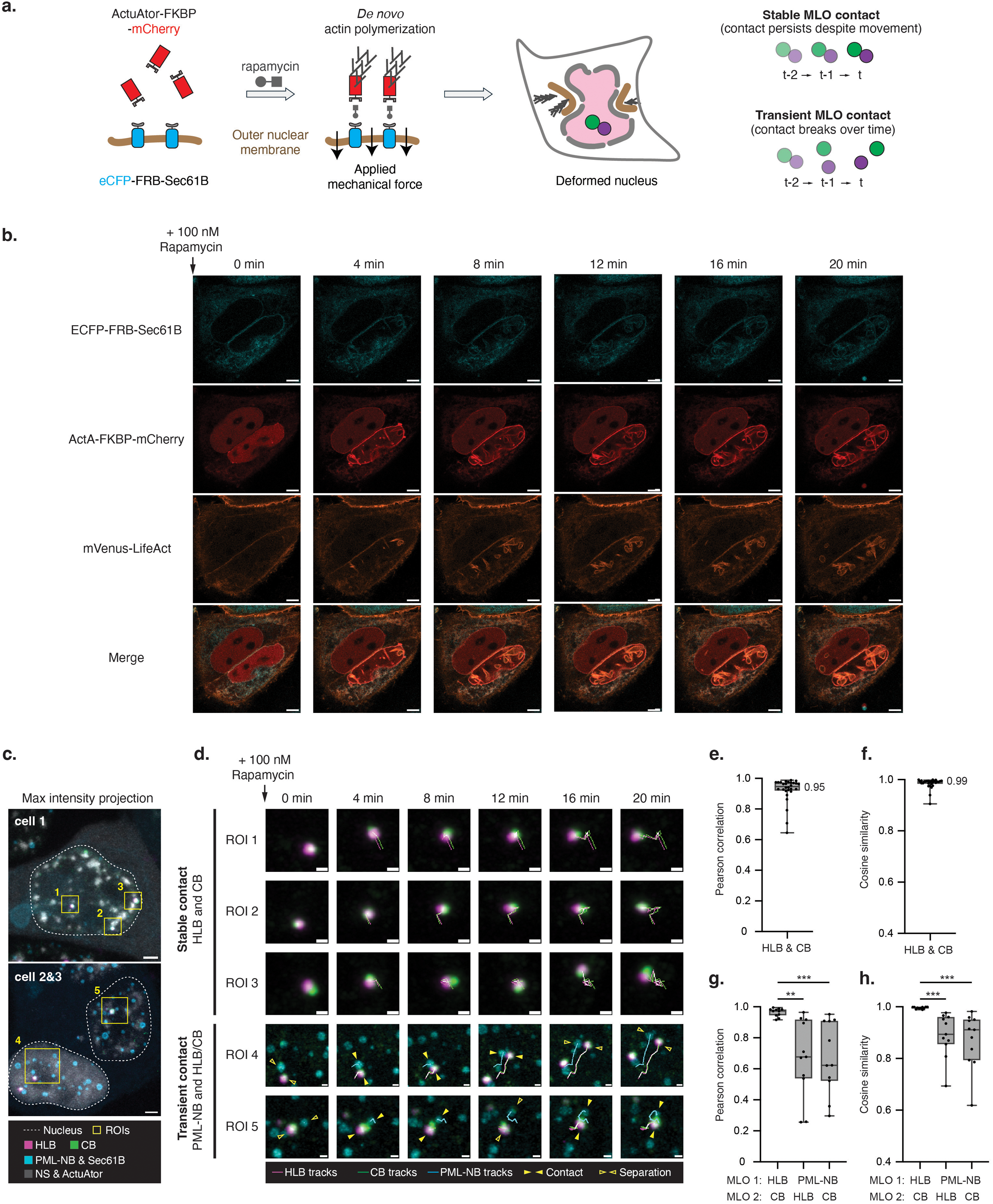
Actuator-induced nuclear deformation validates differential MLO contact modalities via distinct displacement coordination. **(a)** Schematic figure showing the working mechanism of inducible nuclear deformation using the ActuAtor system. ActuAtor (ActA)-FKBP is recruited to the outer nuclear membrane via rapamycin-mediated dimerization with FRB-Sec61B, triggering localized *de novo* actin polymerization and application of mechanical forces to deform the nucleus (left). Stable and transient contacts between MLOs respond differently to nuclear deformation: stable contacts persist, while transient contacts are readily disrupted (right). **(b)** Time-lapse imaging verification of the ActuAtor system by rapamycin-induced actin assembly and nuclear deformation. Representative time-lapse images of live HeLa cells expressing ECFP-FRB-Sec61B (cyan), ActA-FKBP-mCherry (red), and mVenus-LifeAct (orange). The addition of rapamycin (100 nM) induces recruitment of ActA to the nuclear envelope via FRB-FKBP interaction. This recruitment induces actin polymerization at the nuclear periphery, as detected in the mVenus-LifeAct channel, and leads to progressive nuclear deformation over time. Scale bars, 5 µm. **(c-d)** Multispectral 3D time-lapse imaging reveals MLO pair dynamics under nuclear deformation. (**c**) Representative maximum intensity projections of “Rainbow Nucleus” cells transiently expressing ActA-FKBP-mCherry and eCFP-FRB-Sec61B at the initial time point of time-lapse imaging. Scale bars, 2 µm. (**d**) Magnified time-lapse images of the boxed ROIs in (**c**), showing representative displacement tracks of MLO pairs (HLB and CB, ROIs 1-3; PML-NB and HLB or CB, ROIs 4-5) under nuclear deformation following rapamycin treatment (100 nM). Scale bars, 0.5 µm. **(e-f)** Box-and-whisker plots showing the track displacement (**e**) and stepwise directional (**f**) correlations of HLB-CB pairs, indicating highly coordinated movement between HLB and CB under nuclear deformation. For each MLO, centroid positions were determined at each time point from segmented masks for track correlation calculation. Displacement correlation was quantified using Pearson correlation of step displacements between HLB and CB, and directional correlation was quantified as the median cosine similarity of stepwise displacement vectors. Sample sizes, n = 28 HLB and CB pairs from 16 live “Rainbow Nucleus” cells. **(g-h)** Box-and-whisker plots showing the displacement (**g**) and directional (**h**) correlations across PML-NB, HLB and CB, indicating reduced coordination between PML-NB tracks and HLB or CB tracks under nuclear deformation. Displacement and directional correlations were calculated using the same method as in (**e-f**). Statistical significance was assessed using two-tailed t-test. Significance levels were annotated as follows: 0.001 < p ≤ 0.01, **; 0.0001 < p ≤ 0.001, ***. Sample sizes, n = 11 triplets of PML-NB, HLB and CB from 10 live “Rainbow Nucleus” cells.

### Stable CB-NS contact facilitates snRNP diffusion

Among the stable interactions “*Rainbow Nucleus*” uncovered, those between the CB and NS have not been reported or characterized before. CBs are the key assembly and maturation hubs for spliceosomal small ribonucleoproteins (snRNPs), whereas NSs accumulate snRNPs and splicing factors and are critical in the storage and recycling of the splicing machinery^1,8^. We therefore hypothesize that stable CB-NS contacts promote snRNP trafficking between CBs and NSs. To test this, we used single particle tracking (SPT) to assess the diffusion of SmD1, a core Sm-ring protein bound stably to snRNAs during their nuclear transit^23^, in HeLa cells harboring coilin-mStayGold and mScarlet3-SRRM2 knock-ins (**Fig. 6a**). We first confirmed that the Halo-tagged *SNRPD1* (encoding SmD1) correctly localized to both CBs and NSs in cells (**Figs. S6a-c**) similar to WT Sm proteins^24–26^. Fluorescently tagged Sm proteins have previously been demonstrated to incorporate into mature snRNPs^27^. To verify Halo tagged SmD1 behaved similarly in our system, we performed RNA immunoprecipitation (RIP)-qPCR and found that SmD1-Halo enriched substantially more U1, U2, U4, and U5 snRNAs than the NLS (nuclear localization sequence)-Halo control (**Fig. S6d**). As expected, SmD1-Halo did not enrich U6 snRNA which primarily binds Lsm proteins^28–30^ (**Fig. S6d**).

**Figure 6.**
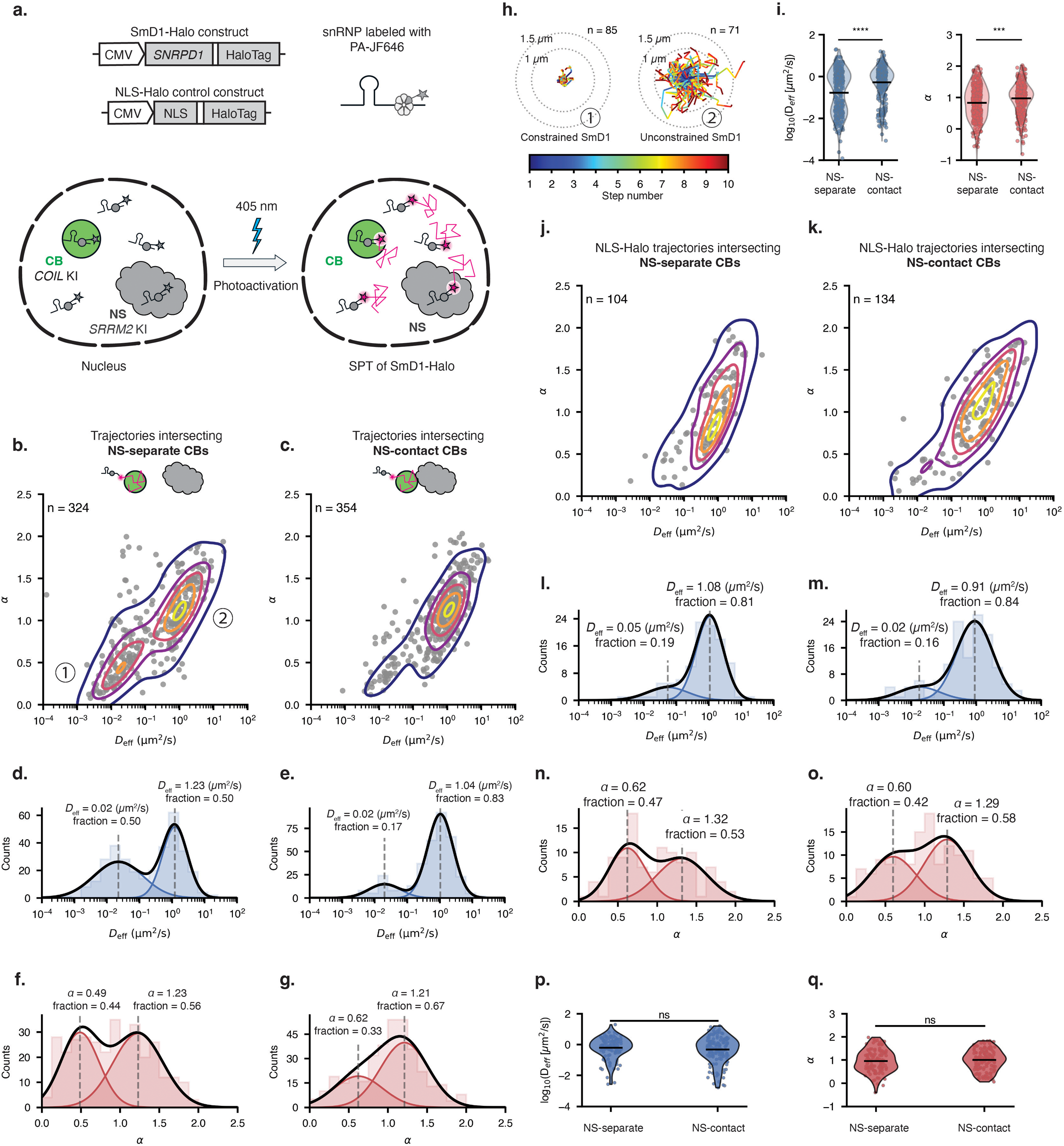
CB contact states with NS are associated with distinct diffusive dynamics of cargo snRNPs. **(a)** Schematic figure demonstrating the workflow of single-particle tracking (SPT) to monitor snRNP dynamics in relation to CB-NS contact states. SmD1-Halo was sparsely photoactivated and tracked in cells containing endogenous CB and NS fluorescent knock-in markers. NLS-Halo served as a control. **(b-c)** 2D probability density plots showing the diffusion states of SmD1-Halo for trajectories intersecting NS-separate CBs (**b**) and NS-contact CBs (**c**). Each point represents a single trajectory plotted by anomalous coefficient (α) versus effective diffusion coefficient (*D*_eff_). Contour lines represent regions of equal probability density regions corresponding to 10%, 35%, 55%, 75%, and 95% of maximum density. Clusters labeled Ill and Ill in (**b**) denote distinct diffusion populations based on α and *D*_eff_. Sample sizes are n = 324 tracks collected from 13 live cells in (**b**), and n = 354 tracks collected from 17 live cells in (**c**). **(d-e)** Histogram plots showing the distribution of *D*_eff_ for SmD1-Halo trajectories intersecting NS-separate CBs (**d**) and NS-contact CBs (**e**). Histograms represent trajectory counts using 20 logarithmically spaced bins. Distributions are fit with two-component Gaussian mixture (GMM) models in log10 scale. Colored curves indicate individual Gaussian components, and the black curve represents the combined model. Dashed lines mark the mean *D*_eff_ of each component, with corresponding proportions labeled. **(f-g)** Histogram plots showing the distribution of α for SmD1-Halo trajectories intersecting NS-separate CBs (**f**) and NS-contact CBs (**g**). Histograms represent trajectory counts using 20 linearly spaced bins. Distributions are fit with two-component GMM models in linear scale. Colored curves indicate individual Gaussian components, and the black curve represents the combined model. Dashed lines mark the mean α of each component, with corresponding proportions labeled. **(h)** Representative trajectories of SmD1-Halo corresponding to the constrained Ill and unconstrained Ill diffusion populations identified in (**b**). Trajectories are colored by step numbers and overlaid with concentric dashed circles indicating 1 µm and 1.5 µm radii. **(i)** Violin plots showing the distributions of *D*_eff_ (left) and α (right) for SmD1-Halo trajectories intersecting NS-separate and NS-contact CBs. Each point represents an individual trajectory. n values are the same as in (**b-c**). Black horizontal lines denote mean values. Statistical significance was assessed using Mann-Whitney U test. Significance levels were annotated as follows: 0.0001 < p ≤ 0.001, ***; p ≤ 0.0001, \*\***\*\***. **(j-k)** 2D probability density plots showing the diffusion states of NLS-Halo for trajectories intersecting NS-separate CBs (**j**) and NS-contact CBs (**k**). Each point represents a single trajectory plotted by anomalous coefficient (α) versus effective diffusion coefficient (*D*_eff_). Contour lines represent regions of equal probability density regions corresponding to 10%, 35%, 55%, 75%, and 95% of maximum density. Sample sizes are n = 104 tracks collected from 8 live cells in (**j**), and n = 134 tracks collected from 10 live cells in (**k**). **(l-m)** Histogram plots showing the distribution of *D*_eff_ for NLS-Halo trajectories intersecting NS-separate CBs (**l**) and NS-contact CBs (**m**). Histograms represent trajectory counts using 20 logarithmically spaced bins. Distributions are fit with two-component Gaussian mixture (GMM) models in log10 scale. Colored curves indicate individual Gaussian components, and the black curve represents the combined model. Dashed lines mark the mean *D*_eff_ of each component, with corresponding proportions labeled. **(n-o)** Histogram plots showing the distribution of α for NLS-Halo trajectories intersecting NS-separate CBs (**n**) and NS-contact CBs (**o**). Histograms represent trajectory counts using 20 linearly spaced bins. Distributions are fit with two-component GMM models in linear scale. Colored curves indicate individual Gaussian components, and the black curve represents the combined model. Dashed lines mark the mean α of each component, with corresponding proportions labeled. **(p-q)** Violin plots showing the distributions of *D*_eff_ (**p**) and α (**q**) for NLS-Halo trajectories intersecting NS-separate and NS-contact CBs. Each point represents an individual trajectory. n values are the same as in (**j-k**). Black horizontal lines denote mean values. Statistical significance was assessed using Mann-Whitney U test. Significance levels were annotated as follows: p > 0.05, ns, indicating no significant difference in the diffusive states of NLS-Halo between NS-contact and NS-separate CBs.

After confirming that SmD1-Halo is functional inside cells, we labeled it with a PA (photo-activatable)-JF646 dye^31^ and used low-level 405 nm laser to sparsely photoactivate individual SmD1 molecules for SPT. Trajectories intersecting CBs were subsequently identified and analyzed to quantify snRNP dynamics (**Fig. 6a**). To understand the roles of CB-NS contacts in snRNP diffusion, we divided SmD1-Halo tracks into two categories: those originating from NS-separate CBs and those from NS-contact CBs. We plotted the distributions of effective diffusion coefficient (*D*_eff_) and anomalous diffusion exponent (α) (**Figs. 6b-c**) and applied two-component Gaussian mixture modeling to identify distinct diffusion states (**Figs. 6d-g**). SmD1-Halo trajectories in NS-separate CBs segregate into two populations: a slow, constrained population (*D*_eff_ = 0.02 µm^2^/s, α = 0.49) and a fast, unconstrained population (*D*_eff_ = 1.23 µm^2^/s, α = 1.23), with approximately equal proportions (**Figs. 6b, d, f**). The constrained tracks were confined within CB-sized regions, whereas the unconstrained tracks explore a larger area extending beyond CB boundaries (**Fig. 6h**), suggesting that during the observation window around half of SmD1-Halo cannot freely diffuse out of CB when CB is not contacting NS. In contrast, SmD1-Halo from NS-contact CBs significantly increased the fraction of mobile, unconstrained tracks (high-*D*_eff_ fraction from 0.50 to 0.83; high-α fraction from 0.56 to 0.67), leading to higher overall *D*_eff_ and α (**Figs. 6c,e,g,i**). As a negative control, we performed the same SPT and analysis using NLS-Halo. In contrast to SmD1-Halo, NLS-Halo exhibited nearly identical diffusion behaviors in NS-separate and NS-contact CBs, showing similar *D*_eff_ and α distributions (**Figs. 6j-k**), diffusion state fractions (**Figs. 6l-o**) and average values (**Figs. 6p-q**), indicating that the change in diffusion dynamics is specific to snRNP components rather than a general consequence of CB-NS proximity. Together, these results indicate that CB-NS contact promotes the trafficking of snRNPs out of CBs.

### RNA Polymerase II inhibition rewires the MLO interactome

Having established that nuclear MLOs organize into a structured, asymmetric, and functional interactome under homeostasis, we next sought to identify the underlying molecular forces that maintain this global spatial organization. Because RNAs play essential roles in organizing various MLOs^32–34^, we hypothesize that active transcription facilitates the interactions among different MLOs. To test this, we selectively perturbed RNA polymerases (Pol) I, II, and III using CX-5461, triptolide, and ML-60218, respectively, and visualized the resulting changes in nuclear MLO organization using the “*Rainbow Nucleus*” (**Fig. 7a**). We then quantified the resulting changes in the interactome (**Figs. 7b-d**), spatial organization (**Figs. 7e-g**) and contact frequencies compared to random levels (**Figs. 7h-j**). We found that different polymerases play distinct roles in organizing the nuclear architecture.

**Figure 7.**
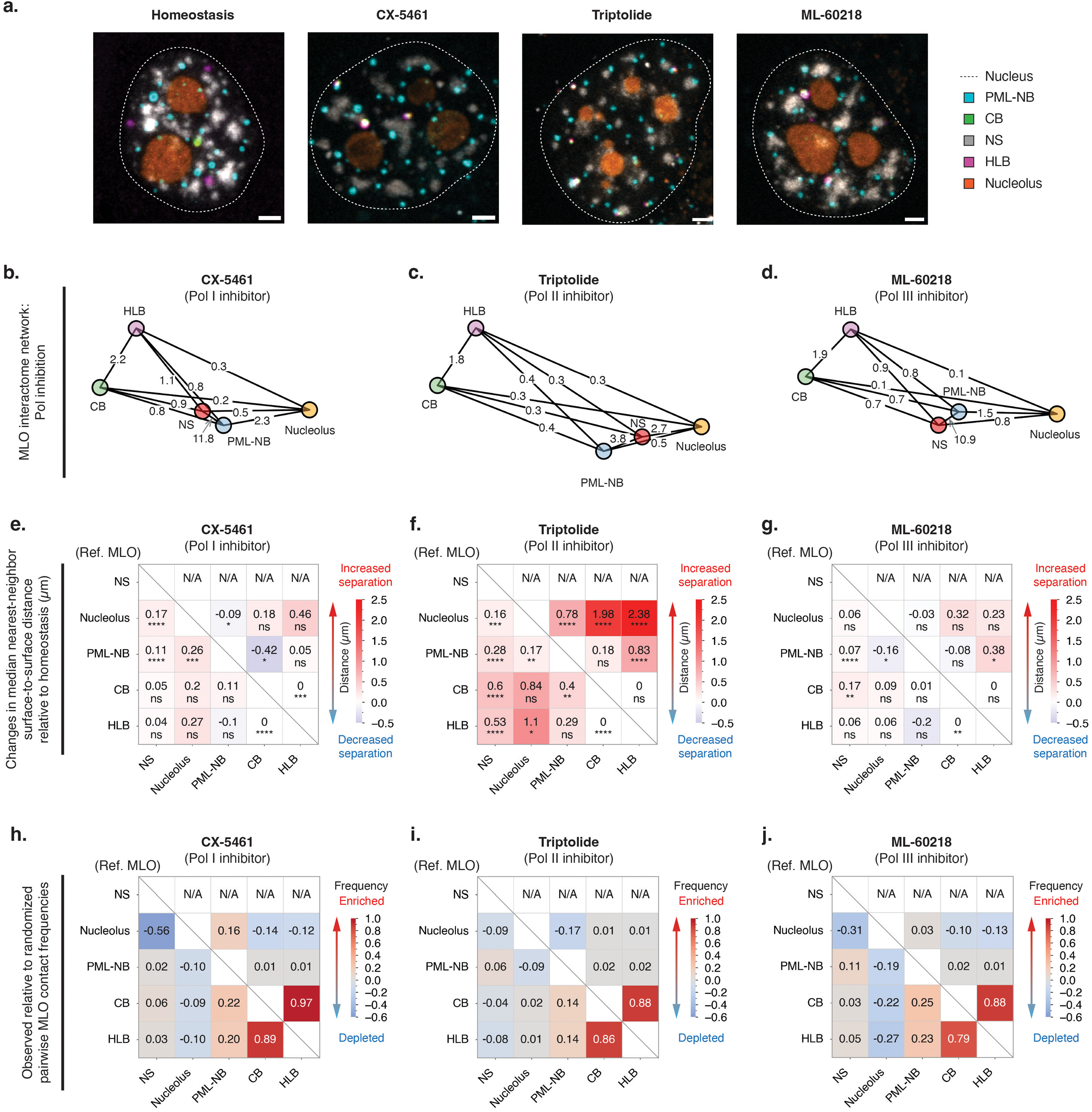
Pol II inhibition induces extensive reorganization of the nuclear MLO interactome. **(a)** Representative maximum intensity projection images of “Rainbow Nucleus” cells showing nuclear MLO organization under homeostasis and transcriptional inhibition conditions. Cells were treated with CX-5461 (0.5 μM), triptolide (0.5 μM), or ML-60218 (40 μM) for 3 hours. Scale bars, 2 µm. **(b-d)** Node-edge network diagrams illustrating pairwise interactions among nuclear MLOs under transcriptional inhibition conditions. Cells were treated with CX-5461 (0.5 μM), triptolide (0.5 μM), or ML-60218 (40 μM) for 3 hours. n = 33 fixed “Rainbow Nucleus” cells for CX-5461 treatment, n = 30 fixed “Rainbow Nucleus” cells for triptolide treatment, and n = 30 fixed “Rainbow Nucleus” cells for ML-60218 treatment. **(e-g)** Heatmaps showing changes in median nearest-neighbor surface-to-surface distances relative to homeostasis under transcriptional inhibition conditions in “Rainbow Nucleus” cells. Cells were treated with CX-5461 (0.5 μM), triptolide (0.5 μM), or ML-60218 (40 μM) for 3 hours. Color indicates the change in median distance between each reference MLO (row labels, left) and target MLO (column labels, bottom), with increased separation (red) and decreased separation (blue). Sample sizes are n = 82 (HLB), 75 (CB), 1198 (PML-NB), and 89 (nucleolus), collected from 33 fixed “Rainbow Nucleus” cells for CX-5461 treatment; n = 61 (HLB), 60 (CB), 495 (PML-NB), and 210 (nucleolus) from 30 fixed “Rainbow Nucleus” cells for triptolide treatment; and n = 70 (HLB), 63 (CB), 890 (PML-NB), and 74 (nucleolus) from 30 fixed “Rainbow Nucleus” cells for ML-60218 treatment. Statistical significance was assessed using two-tailed Mann-Whitney U tests comparing each transcriptional inhibition condition to homeostasis controls. Significance levels were annotated as follows: p > 0.05, ns; 0.01 < p ≤ 0.05, *; 0.001 < p ≤ 0.01, **; 0.0001 < p ≤ 0.001, ***; p ≤ 0.0001, \*\***\*\***. **(h-j)** Heatmaps showing observed relative to randomized pairwise MLO contact frequencies among nuclear MLOs in “Rainbow Nucleus” cells treated with CX-5461 (0.5 μM, left), triptolide (0.5 μM, middle), or ML-60218 (40 μM, right) for 3 hours. Color indicates the difference between observed and randomized contact frequencies for each pairwise MLO combination, with enrichment (red) and depletion (blue) relative to random expectation. Sample sizes are the same as in (**e-g**). For each treatment condition, randomization analysis was performed using segmented MLO masks derived from 10 drug-treated fixed “Rainbow Nucleus” cells.

Inhibition of Pol I transcription via a 3-hour treatment with 0.5 µM CX-5461 effectively depleted nascent RNAs within the nucleolar region as visualized by 5-ethynyl uridine (EU) staining (**Figs. S7a-b**). However, the overall architecture of the MLO interactome remained largely unchanged (**Fig. 7b**). MLO spatial organization exhibited only minor deviations from homeostasis, characterized by slight increases in the separation of all MLOs from NSs—particularly for the nucleolus–NS and PML-NB–NS pairs—alongside closer proximity between PML-NB–CB and PML-NB–HLB (**Figs. 7e, S7c, S8a**). This implies that while localized ribosomal RNA (rRNA) synthesis by RNA Pol I acts as an internal nucleolar scaffold^34,35^, RNA Pol I activities do not mediate long-range inter-MLO interactions outside the nucleolus. Similarly, Pol III inhibition by ML-60218 produced mild effects on the nuclear MLO interactome. Compared with homeostasis, the overall interactome architecture was largely preserved (**Figs. 7d, S7c, S8a**), with only minor increases in separation between NS and other MLOs alongside selective decreases in distances between PML-NB and the nucleolus (**Fig. 7g**). This minimal disruption implies that short, highly structured Pol III transcripts (such as tRNAs) do not contribute significantly to the interactions between MLOs. Consistent with these observations, following Pol I or Pol III inhibition, MLO contact frequencies compared to random levels remained largely similar to homeostatic levels (**Fig. 3d)**, with the primary changes being reduced enrichment of NS-associated contacts (**Figs. 7h,j, first column, S8b-c**), arising from the increase in separation between NS and other MLOs. Together, these perturbations demonstrate that Pol I and Pol III transcription, despite their roles in specific biosynthetic processes, do not maintain the global MLO interactome.

In contrast, Pol II inhibition by triptolide induced extensive rewiring of the nuclear MLO interactome. Treatment with 0.5 µM triptolide for 3 hours depleted nascent RNAs in the nucleoplasm (**Figs. S7a-b**), reduced global MLO contact counts (**Fig. 7c**), and significantly increased pairwise MLO surface-to-surface distances, resulting in a pronounced separation of MLOs from each other (**Figs. 7f, S7c, S8a**). Accompanying this spatial dissociation, the structured, asymmetric MLO interactome largely collapsed: except for the CB–HLB pair, most pairwise MLO contact frequencies became indistinguishable from randomized associations (**Figs. 7i, S8b-c**). These results suggest that Pol II–dependent transcripts may act as multivalent bridges linking MLOs, although the present data cannot exclude indirect consequences of transcriptional arrest. Consequently, active Pol II transcription—rather than the general transcriptional activity—is uniquely and broadly required to prevent the higher-order nuclear MLO interactome from collapsing into a near-random organizational state.

## Discussion

Our study presents the first comprehensive, live-cell interaction map of five major nuclear MLOs imaged simultaneously. Through multi-spectral, live-cell imaging using “*Rainbow Nucleus*,” we uncovered specific interaction modes among different MLOs. Spatially, NS acts as a central interaction hub, whereas the nucleolus remains largely isolated from other MLOs. Temporally, the HLB-CB, CB-NS, and HLB-NS pairs form stable contacts, while the remaining MLO pairs engage in transient or no interactions. These results raise new questions regarding inter-MLO dynamics, such as what mediates these stable versus transient states and what functions they serve.

Our data shows that transcription, specifically that mediated by RNA Pol II, is essential in sustaining the global MLO interactome. The products of RNA Pol II, messenger RNAs and long non-coding RNAs regulate the formation and material properties of different MLOs^36–38^, and may also mediate contacts among different MLOs. Alternatively, protein tethers, like those linking MBOs^39^ may mediate stable contacts among different MLOs. These RNA and protein tethers can be identified in the future by finding overlaps of different MLO-specific RNA and proteomics datasets^40,41^, or using the Split-APEX2 and/or Split-TurboID which are especially suited for identifying RNA/proteins at the interface^42–44^. Follow-up studies using super resolution imaging or correlative light and electron microscopy (CLEM) are needed to verify these tethers. On the other hand, interactions may also occur based on opposing molecular charges of MLOs dictated by pH^5^. Highly sensitive intranuclear pH dyes will be useful to test this theory. Small nuclear RNAs, which are also products of RNA Pol II, can flow between different MLOs and in turn bring into proximity of these MLOs. Experiments and theories need to be combined to test this hypothesis.

In addition to identifying the specific proteins/RNAs mediating these contacts, what is more pivotal is to understand the functions of these MLOs interactions. Our study shows that one of the stable contacts we discovered, the CB-NS contact, facilitate the diffusion of snRNPs. Nature is full of examples of transferring materials via direct contacts: cytonemes are specialized nanotubes formed in developing tissues to transfer hedgehog molecules from one cell to another^45^; membrane contact sites between ER and mitochondria facilitate the transfer of lipids and Ca^2+^ between these organelles^39^. While it is unclear why direct contacts between CBs and NSs are needed for snRNP transfer, we propose that the specific conformation of the assembled snRNPs can be maintained through direct transfer when they do not need to contact the nucleoplasm^8^. On the other hand, transient inter-MLO interactions may mediate post-translational modifications such as SUMOylation mediated by the kissing of PML-NBs. Future studies of these stable and transient interactions—and the development of tools to modulate them—may ultimately enable us to fine-tune nuclear organization for regenerative medicine and disease therapies.

## Limitations of the study

We understand that the FRAP assay using one marker protein of each MLO cannot fully reveal the material properties of each MLO. Future studies are needed to probe additional components of the MLO, using methods such as single particle tracking and *in vitro* reconstitutions.

## Resource availability

### Lead contact

Further information and requests for resources and reagents should be directed to and will be fulfilled by the lead contact Danfeng Cai.

### Materials availability

All materials generated in this study are available upon request.

### Data and code availability

- Custom image analysis python codes are available in Github: https://github.com/BMBCaiLab/2026-Rainbow-Nucleus-paper.git
- Any additional information required to reanalyze the data reported in this paper is available from the lead contact upon request.

## Supporting information

Key Resources Table

## Acknowledgements

We appreciate the feedback and discussions from Sarah Cohen (UNC), Paul Tillberg (HHMI-Janelia Research Campus), and Cai lab members. We are grateful to Henrietta Lacks, now deceased, from whom HeLa cell line was established without her knowledge or consent in 1951, which made significant contributions to scientific progress and advances in human health. We also acknowledge the Johns Hopkins University (JHU) Microscope Facility for access to confocal microscopy, supported by NIH grant S10OD023548 (Scot Kuo). This study is supported by the National Institutes of Health (R35GM142837, D.C., R35GM149329, T.I.), the Kleberg Foundation (T.I.), Johns Hopkins Catalyst Award (D.C.), and Johns Hopkins Provost’s Undergraduate Research Award (N.B.G.).

## Author contributions

Conceptualization: D.C.; Methodology and investigation: S.Y., N.B.G., C.L.S.; Resources: T.I.; Software: S.Y.; Supervision: D.C.; Visualization: S.Y.; Writing – original draft: D.C.; Writing – review & editing: S.Y., D.C.; Funding acquisition: T.I., D.C.

## Supplemental information titles and legends

Document S1. Figures S1-S8, Table S1

**Movie S1. Global dynamics of nuclear membrane-less organelles (MLOs) in the “Rainbow Nucleus” cell nucleus**.

Representative movie of a live “Rainbow Nucleus” cell nuclear region to simultaneously visualize the dynamics of five nuclear MLOs, including histone locus bodies (HLBs, magenta), Cajal bodies (CBs, green), nuclear speckles (NSs, red), PML nuclear bodies (PML-NBs, cyan), and nucleoli (orange). Smaller panels on the right indicated the individual fluorescence channels corresponding to each nuclear MLO. Scale bar, 2 µm.

**Movie S2. Stable HLB-CB association with transient “kiss-and-run” contacts from a neighboring PML-NB**.

Representative movie of a magnified region of a live “Rainbow Nucleus” cell showing a stable HLB (magenta)-CB (green) pair accompanied by transient “kiss-and-run” interactions from a neighboring PML-NB (cyan). While the HLB-CB pair remains continuously associated throughout the imaging period, PML-NB exhibit transient and short-lived contact events with the HLB and CB. Scale bar, 1 µm.

**Movie S3. HLB and CB maintain stable contacts with NS**.

Representative movie of a magnified region in a live “Rainbow Nucleus” cell showing a stable contact between HLB (magenta)-CB (green) pair and the surrounding NS (red). Scale bar, 1 µm.

**Movie S4. PML-NB exhibit transient contacts with NS**.

Representative movie of a magnified region in a live “Rainbow Nucleus” cell showing transient contacts between a PML-NB (cyan) and the surrounding NS (red). Scale bar, 1 µm.

**Movie S5. *In situ* nuclear actin formation enabled by molecular tool Actuator.** Representative movie of HeLa cells showing rapid formation of nuclear actin structures following actuator activation by addition of 100 nM rapamycin. mCherry-FKBP-ActA is shown in red, mVenus-LifeAct in orange, and ECFP-FRB-Sec61B in cyan. Smaller panels on the right indicated the individual fluorescence channels. Scale bar, 2 µm.

**Movie S6. Actuator-induced actin formation disrupts nuclear MLO positions.**

Representative movie of “Rainbow Nucleus” cells showing changes in the positioning and movement of nuclear MLOs following actuator-induced nuclear actin assembly. HLB is shown in magenta, NS and mCherry-FKBP-ActA in red, CB in green, and PML-NB and ECFP-FRB-Sec61B in cyan. Cells transfected with the actuator system (center 2 cells) exhibit pronounced repositioning and dynamic rearrangement of nuclear MLOs, whereas the non-transfected cell (bottom right) displays comparatively stable MLO positioning with minimal spatial changes over time. Smaller panels on the right indicated the individual fluorescence channels. Scale bar, 2 µm.

## Materials and Methods

### 1. Plasmids construction

#### Coilin gRNA-Cas9 and SRRM2 gRNA-Cas9 plasmid construction

The guide RNAs targeting the C-terminus of *COIL* locus (5′-gagattcaataatcagtctt-3′) and the guide RNA targeting the N-terminus of *SRRM2* locus (5’-gggccatgtacaacgggatc-3’) were cloned into pSpCas9(BB)-2A-GFP vector following the scarless cloning method described by Ran *et al*^46^. The pSpCas9(BB)-2A-GFP (pX458) was a gift from Feng Zhang Lab (Addgene plasmid # 48138; http://n2t.net/addgene:48138; RRID:Addgene_48138). Target-specific guide sequences containing overhangs compatible with pSpCas9(BB) BbsI sites were ordered (Integrated DNA Technologies, IDT), then annealed and phosphorylated in a 10 µL reaction containing 1 µL 100 µM forward oligo, 1 µL 100 µM reverse oligo, 1 µL 10x T4 ligation buffer (B0202S, NEB), 1 µL T4 polynucleotide kinase (B0201S, NEB), and 6 µL nuclease-free water. The mixtures were incubated at 37°C for 30 min, heated to 95°C for 5 min, then cooled to 25°C at a ramp rate of – 5°C/min in a thermocycler (C1000 Touch^TM^, Bio-Rad). The annealed oligos were ligated into pX458 vector in a single-tube digestion-ligation reaction containing 100 ng pX458 vector, 1 µL 100 nM oligo duplex, 1 µL FastDigest BpiI (FD1014, ThermoFisher), 2 µL 10x FastDigest buffer, 1 µL T4 DNA ligase (M0202S, NEB), and nuclease-free water to a final volume of 20 µL. The reactions were incubated at 37°C for 1 hour, followed by alternating 5 min cycles at 37°C and 21°C for six rounds. The ligations were transformed into DH5α competent cells (EC0112, ThermoFisher), individual clones were isolated and amplified. The correct inserts were verified by Sanger sequencing.

#### Coilin and SRRM2 donor repair plasmid construction

The donor repair plasmids were constructed in pUC57-Mini vector (GenScript). Gene fragments encoding the 5’ homology arm, in-frame linker, fluorescent protein sequence, and the 3’ homology arm for *COIL* and *SRRM2* genes were commercially synthesized. mStayGold^47^ was used as fluorescent proteins tags for *COIL* and mScarlet3^48^ was used for *SRRM2* Empty. pUC57-Mini vector was linearized by EcoRV-HF^®^ (R3195S, NEB) and further purified (QIAquick PCR Purification Kit, 28104, QIAGEN) prior to ligation. The synthetic gene fragments and linearized empty vector were assembled using NEBuilder HiFi (E2621L, NEB) at 50°C for 15 minutes. The ligations were transformed into DH5α competent cells (EC0112, ThermoFisher), individual clones were isolated and amplified. The correct inserts were verified by Sanger sequencing.

#### BFP-hPMLIII and Halo-mini-FLASH lentiviral plasmid construction

Both BFP-hPMLIII and Halo-mini-FLASH lentiviral transfer plasmid were constructed in pLVX-Puro vector (Takara Bio). To prevent overexpression of the protein of interest, which causes the abnormal size of target membrane-less organelles, the original CMV promoter in pLVX-Puro was replaced with a moderate-activity polyubiquitin C promoter. The coding sequence for BFP-hPMLIII was derived from pCIBN-tagBFP-hPMLIII (Addgene #103807, a gift from Karsten Rippe Lab), and the mini-FLASH coding sequence was a gift from Michael Matunis Lab. Mini-FLASH is a fusion of amino acids 1-200 and 1751-1982 of the full-length FLASH and has been shown to be recruited to the histone locus body (HLB) as efficiently as the WT FLASH protein reference^49,50^. Gene fragments encoding the desired constructs were PCR-amplified from the respective source plasmids, ligated with linearized pLVX-Puro backbone using the NEBuilder HiFi DNA Assembly Kit (E2621L, NEB). The ligations were transformed into DH5α competent cells (EC0112, ThermoFisher), and verified by whole plasmid sequencing (Plasmidsaurus Inc.).

#### LSS-mKate 2-NPM1 plasmid construction

LSS-mKate2-NPM1 was constructed in a pCMV vector. Briefly, gene insert encoding the LSS-mKate2^51^, a flexible linker and NPM1 was commercially synthesized, and flanked with homologous sequences at both ends for assembly with the linearized pCMV backbone. The ligation was conducted with NEBuilder HiFi DNA Assembly Kit (E2621L, NEB), transformed into DH5α competent cells (EC0112, ThermoFisher), and verified by whole plasmid sequencing (Plasmidsaurus Inc.).

### 2. Cell cultures

All cell lines used in this study were cultured in Dulbecco’s Modified Eagle’s Medium (DMEM, 15-013-CV, Corning) supplemented with 10% v/v fetal bovine serum (26140079, Gibco), 1% v/v GlutaMAX^TM^ supplement (35050061, Gibco) and 1% v/v Penicillin-Streptomycin (10,000 U/mL) (15140122, Gibco). Cell cultures were incubated at 37°C with 5% CO_2_ in a humidified incubator. For live-cell imaging experiments, the culture medium was changed to FluoroBrite^TM^ DMEM (A1896701, Gibco) supplemented with 10% v/v fetal bovine serum (26140079, Gibco), 1% v/v GlutaMAX^TM^ supplement (35050061, Gibco) and 1% v/v Penicillin-Streptomycin (10,000 U/mL) (15140122, Gibco).

### 3. Cell lines

#### HEK293T cells

HEK293T cells were acquired from ATCC (CRL3216).

#### HeLa cells

Hela cells were acquired from ATCC (CCL-2). The HeLa cell line was further authenticated through Short Tandem Repeat (STR) profiling.

#### “*Rainbow Nucleus*” cell line generation

“*Rainbow Nucleus*” cells were genetically engineered to express marker proteins for various membrane-less organelle (MLO), each tagged with distinct fluorescent proteins or a HaloTag self-labeling protein tag for multiplexed imaging. The “*Rainbow Nucleus*” cell line was generated using a mixed strategy. First, fluorescent proteins mStayGold and mScarlet3 genes were inserted in *COIL* and *SRRM2* locus sequentially by CRISPR mediated knock-in strategy. Next, to further establish stable cell lines expressing TagBFP-PML and Halo-mini-Flash, lentiviral transductions were performed using second-generation lentiviral systems. Cells were subjected to fluorescence-activated cell sorting (FACS) after each round of stable cell line generation to isolate single cell clones. Isolated cell lines were further authenticated via genomic PCR, western blotting and immunofluorescence imaging. Finally, a plasmid encoding LSS-mKate 2-NPM1 was transiently transfected into the established cell line to label the nucleolus 16 hrs prior to imaging. The detailed procedures of each round of stable cell line generation were outlined below.

##### HeLa Coilin-mStayGold Cell Line Generation

Endogenously tagged HeLa Coilin-mStayGold cells were engineered using CRISPR-Cas9 system. HeLa cells were seeded in a 35 mm glass bottom imaging dish (D35-20-1.5-N, Cellvis) one day prior to the transfection. Upon confluency reaches ∼70% in the following day, cells were co-transfected with a sgRNA-Cas9-eGFP pSpCas9 expression plasmid and mStayGold-encoding repair DNA fragments with Lipofectamine 3000 Transfection Regent (L3000015, ThermoFisher) following the manufacturer’s protocol. Transfection efficiency was assessed 24 hours post-transfection by monitoring Cas9-eGFP fluorescence. At 48 hours post-transfection, cells were passaged to a T-25 flask for expansion. FACS was performed one week after initial transfection to isolate HeLa Coilin-mStayGold cell lines, allowing sufficient time for transient Cas9-eGFP to subside and avoid interference with Coilin-mStayGold signal. Isolated single-cell colonies were allowed to expand in 96-well plates for 3 weeks. Genomic DNA was extracted from expanded clones using Quick-DNA^TM^ MicroPrep Kit (D3020, Zymo Research), and targeted modified gene locus was amplified by PCR using Q5 High-Fidelity 2x Master Mix (M0492, NEB). PCR products were analyzed by agarose gel electrophoresis to confirm the expected amplicon size further validated by sequencing. The resulting HeLa Coilin-mStayGold cell line was further validated by immunofluorescence imaging and western blotting.

##### HeLa Coilin-mStayGold, mScarlet3-SRRM2 Cell Line Generation

Based on the established HeLa Coilin-mStayGold cell line, a dual-tagged HeLa Coilin-mStayGold::mScarlet3-SRRM2 cell line was generated using the same CRISPR-Cas9 mediated knock-in strategy. A PX458-based Cas9 plasmid encoding sgRNA targeting 5’ region of the human *SRRM2* locus was co-transfected with a linearized donor DNA block encoding a mScarlet3 fusion with homology arms. FACS was performed one-week post-transfection to isolate mScarlet3-positive clones. The resulting HeLa Coilin-mStayGold, mScarlet3-SRRM2 knock-in cell line was further tested by genomic PCR validation, western blotting and immunofluorescence imaging.

##### HeLa Coilin-mStayGold, mScarlet3-SRRM2, BFP-hPML III Cell Line Generation

To further integrate fluorescent protein markers to label PML-NBs, we generated stably cell lines expressing BFP-hPML III using second-generation lentiviral transduction systems. HEK293T cells were used to generate lentiviral particles. Briefly, HEK293T cells were seeded in T-75 flask one day prior to the transfection. Upon confluency reaching ∼70% in the following day, cells were co-transfected with 10 µg of BFP-hPML III transfer plasmid, 6 µg of packaging plasmid D8.91 and 2 µg of envelope plasmid VSV-G (both D8.91 and VSV-G plasmids are gifts from Utthara Nayer Lab) using Lipofectamine 2000 (11668027, ThermoFisher). The medium was replaced after 24 hours with fresh medium. Supernatant containing lentiviral particles was harvested twice, at 48 and 72 hours post-transfection, and stored at –80 °C for future use. For lentiviral transduction, HeLa Coilin-mStayGold, mScarlet3-SRRM2 knock-in cell was transduced with viral supernatant supplemented with 8 µg/mL polybrene (TR-1003-G, Sigma-Aldrich) for 24 hours. After recovery in fresh medium, cells were harvested 72 hours post-transduction and subjected to FACS. The BFP-positive sub-population was sorted into fresh growth medium for clonal expansion. The resulting cell line was further validated by immunofluorescence imaging and western blotting.

##### HeLa Coilin-mStayGold, mScarlet3-SRRM2, BFP-hPML III, Halo-mini-FLASH Cell Line <u>Generation</u>

Finally, a subsequent round of lentiviral transduction was performed to introduce HaloTag-fused mini-FLASH protein, following the same procedure as described above. This enabled the specific labeling of the histone locus body (HLB) through stable expression of mini-FLASH.

##### LSS-mKate 2-NPM1 transfection

To further mark the nucleolus for live-cell imaging, a total of 2 µg plasmid encoding LSS-mKate2-NPM1 was transiently transfected into the established “*Rainbow Nucleus*” cell line using Lipofectamine 3000 (L3000015, ThermoFisher). Cells were seeded in 35 mm glass-bottom dishes (D35-20-1.5N, Cellvis) one day prior to transfection and reached approximately 70% confluency at the time of transfection. Cells were incubated for 16 hours post-transfection before subsequent imaging.

### 4. Image acquisition

#### Multi-spectral Airyscan imaging of the “*Rainbow Nucleus*” cell lines

Multi-spectral 3D Airyscan imaging was performed on a Zeiss LSM 900 equipped with an Airyscan 2 detector using a Plan-Apochromat 63×/1.4 NA oil immersion objective lens (Carl Zeiss). For HaloTag labeling, 100 nM Janelia Fluor 646 HaloTag ligand^52^ (a gift from Luke Lavis Lab) was added to the culture medium 2 hours prior to imaging. Cells were subsequently washed and maintained in FluoroBrite™ DMEM (A1896701, Gibco) supplemented with 10% fetal bovine serum (26140079, Gibco), 1% GlutaMAX™ (35050061, Gibco), and 1% Penicillin-Streptomycin (15140122, Gibco) before imaging. During imaging, cells were maintained at 37 °C and 5% CO₂ using a stage-top incubation system throughout imaging. Raw Airyscan images were processed using ZEN software (Zeiss) with standard Airyscan deconvolution parameters.

For five-color imaging of the “*Rainbow Nucleus*” cells, sequential 3D acquisition was used to minimize spectral cross-talk. The following excitation lines and emission detection windows were used: 405 nm excitation with 420-485 nm emission collection for TagBFP; 488 nm excitation with 490-530 nm emission collection for mStayGold; 488 nm excitation with 600-700 nm emission collection for LSS-mKate2, 561 nm excitation with 566-650 nm emission collection for mScarlet3; and 640 nm excitation with 645-700 nm emission collection for HaloTag-JF646.

#### 3D Time-lapse imaging for MLO contact dynamics

3D time-lapse imaging was performed using the same sequential multi-track configuration described above. Z-stacks were acquired across the full nuclear volume with a step size of 0.6 µm to balance axial resolution and imaging speed. Time-lapse acquisition was performed at 20 seconds intervals for 10 minutes. Airyscan-processed images stacks were subjected to custom 3D segmentation, object tracking, and interaction analysis pipelines as described below.

#### Actuator-based nuclear deformation to perturb MLO contacts

“*Rainbow Nucleus*” cells were transiently co-transfected with Actuator constructs^22^, 1 µg ActuAtor-FKBP-mCherry and 1 µg eCFP-FRB-Sec61B, in 35 mm glass-bottom dishes (D35-20-1.5N, Cellvis) using Lipofectamine 3000 (L3000015, ThermoFisher). Cells were seeded one day prior to transfection and reached ∼70% confluency at the time of transfection. Following transfection, cells were incubated for 16 h to allow protein expression. For induction of actin polymerization, rapamycin was added at a final concentration of 100 nM to drive FKBP-FRB heterodimerization and recruit the actuator to the outer nuclear membrane. This recruitment triggers *de novo* actin polymerization at the nuclear surface, thereby applying localized mechanical force to the nucleus and inducing nuclear deformation. Following rapamycin treatment, cells were imaged to assess changes of spatial organization and contact behavior of MLOs, using 3D time-lapse imaging as described above.

#### Fluorescence recovery after photobleaching (FRAP)

FRAP experiments were conducted using “*Rainbow Nucleus*” cells prepared in 35 mm glass-bottom dishes (D35-20-1.5N, Cellvis) as described above. FRAP was performed using a Zeiss 900 confocal microscope equipped with 405, 488, 561, and 640 nm lasers. Cells were maintained at 37°C in a humidified stage-top incubator with 5% CO₂ throughout the experiment. Defined region of interest (ROIs) within different MLOs were photobleached using a high-intensity laser pulse, followed by time-lapse acquisition at low laser power to monitor fluorescence recovery. The same imaging settings were applied to monitor unintended photobleaching during acquisition, and the resulting signal loss was accounted for during normalization. Images were collected at 0.5-5 seconds intervals over a total duration of 1-5 minutes, with the settings adjusted to cover the whole recovery curves. Fluorescence recovery curves were normalized by setting the average pre-bleach intensity to 1 and the first post-bleach frame to 0. Mobile fractions were calculated from the plateau value of the fitted recovery curve. Recovery half-times (T_1/2_) were derived using single-exponential fitting in GraphPad Prism.

#### Single particle tracking

Single particle tracking of the SmD1-Halo protein was performed using a Nikon Ti2 Eclipse TIRF microscope equipped with a 100 x/ 1.49 NA oil-immersion objective and a Perfect Focus System to maintain axial stability. Signal was collected via a Hamamatsu ORCA Fusion BT CMOS camera. HeLa Coilin-mStayGold KI, mScarlet3-SRRM2 KI cells were seeded in 35 mm glass-bottom dishes (D35-20-1.5N, Cellvis) one day prior to SmD1-Halo transfection and reached approximately 70% confluency at the time of transfection. Cells were co-transfected with 0.5 µg *SNRPD1-*HaloTag encoding plasmid (synthesized via Twist Bioscience) and 1.5 µg empty pUC57-Mini vector (GenScript) per 35 mm dish for 16 hours with Lipofectamine 3000 Transfection Regent (L3000015, ThermoFisher). Prior to imaging, cells were labeled with 100 nM PA-JF646 HaloTag ligands (a gift from Luke Lavis Lab) for 1 hour diluted in FluoroBrite™ DMEM (A1896701, Gibco) supplemented with 10% fetal bovine serum (26140079, Gibco), 1% GlutaMAX™ (35050061, Gibco), and 1% Penicillin-Streptomycin (15140122, Gibco). Subsequently, cells were washed three times in ligand-free medium at 5 minutes intervals. SPT images was performed under TIRF illumination with an exposure time of 8 ms per frame using 640 nm laser excitation at 100% power (LUN-F XL) at a frame rate of 104.54 frames/s for 5 minutes. A low-intensity 405 nm laser (1% power) was applied concurrently to stochastically and continuously photoactivate PA-JF646, ensuring a sparse density of fluorescent molecules suitable for SPT. Reference images of the same field of view were acquired immediately prior to SPT imaging under TIRF illumination using 488 nm and 560 nm excitation at 100% power (LUN-F XL) with an exposure time of 20 ms per channel. These reference images were used to generate binary masks for CB and NS, respectively. The focal plane and stage position were maintained constant between reference and SPT acquisitions to ensure accurate spatial alignment.

### 5. Image analysis

#### Segmentation of MLOs

Airyscan processed cell images (5D stack, T, Z, C, Y, X) were first exported as 16-bit TIFF files using Fiji^53^ (ImageJ, v2.14.0), and further processed using custom Python scripts to isolate individual channels for downstream contact analysis.

First, for a single time point, image stacks for each channel (3D stack, Z, Y, X) was processed independently. Volumes were first subjected to Gaussian smoothing (σ = 1.5) to reduce pixelated noise. Subsequently, a maximum intensity projection (MIP) image was generated from the smoothed 3D stack, and Otsu thresholding was applied to the normalized MIP to determine a specific mask cutoff intensity for each channel at each timepoint^54,55^. This resulting threshold value was then applied to the corresponding full 3D volume to generate a 3D binary mask. Channel-specific 3D binary masks were further cleaned by removing small objects below certain voxel thresholds (HLB: 20 voxels; NS: 20 voxels; CB: 20 voxels; nucleolus: 3000 voxels; PML-NB: 20 voxels) to exclude non-specific signals and segmentation artifacts. A resulting composite mask dataset (4D, Z, C, Y, X) were constructed and saved for downstream MLO contact analysis.

Next, for multi-timepoint datasets collected from MLO contact dynamics analysis, this above-mentioned thresholding pipeline was applied independently to each timepoint, allowing adaptive threshold determination per timepoints to account for signal variation and photobleaching. Optional user-defined parameters, including fixed-threshold overrides at specific timepoints, were applied when necessary to ensure biological accurate segmentation. Finally, composite mask dataset (5D, T, Z, C, Y, X) were constructed and saved for downstream MLO contact analysis.

#### MLO tracking and contact detection

MLO contact analysis was performed using custom Python scripts on segmented mask datasets to quantify spatial proximity and interaction dynamics among MLOs.

For single timepoint datasets (4D, Z, C, Y, X), contacts were defined based on spatial proximity between segmented MLO objects within the 3D volume. A contact event was defined when the minimum edge-to-edge distance between two MLOs was less than or equal to one pixel (35.4 nm) in the lateral (XY) plane. This threshold was chosen to capture direct physical contact between MLOs while accommodating segmentation variability. Of note, this contact evaluation was performed only in the XY dimensions and not along the Z axis, as the spatial pixel steps (130 nm) in Z is substantially higher than in XY. This constraint minimizes overestimation of contacts due to larger axial steps and lower axial resolution.

For multi-timepoint datasets (5D stacks, T, Z, C, Y, X), object tracking was performed prior to contact analysis. MLOs identified at each timepoint was tracked across consecutive frames using an intersection-over-union (IoU)-based matching strategy^56^. Objects displaying an IoU greater than 0.05 between consecutive timepoints were identified as the same MLO object. This approach enables consistent tracking of individual MLOs despite changes in morphology or position between frames. Tracking results were further inspected to correct potential errors and to manually link objects that exhibited large displacements between frames. Following tracking, contact detection was performed at each timepoint using the same proximity-based criterion described above. For each MLO pairs, consecutive contact timepoints were grouped, and the durations of the contact were recorded. Finally, time-resolved pairwise contacts among five MLO types (HLB, NS, CB, nucleolus, and PML-NB) were summarized in data tables, yielding a total of 10 possible MLO contact combinations.

#### MLO interactome network visualization

Pairwise MLO contacts were quantified from segmented 3D binary masks of MLOs in fixed “*Rainbow Nucleus*” cells. For each cell, the number of contacts between each pair of MLO types was calculated using the method describing above, generating 10 pairwise contact measurements across the five MLO classes. The mean number of pairwise contacts per cell was subsequently calculated for each MLO combination and used to construct the nuclear MLO interactome.

We subsequently used Cytoscape software to generate node-edge network diagrams to illustrate the global interactome among nuclear MLOs similar to Valm *et al*.’s work assessing interactions among membrane-bound organelles (MBOs)^57^. Each node represents one MLO type. Edge labels indicate the mean number of pairwise contacts per cell for each MLO pair. Edge length was manually adjusted to be approximately inversely related to the mean number of contacts per cell, such that MLO pairs with more frequent contacts are positioned closer together, whereas pairs with fewer contacts are positioned farther apart.

#### MLO contact dynamics analysis

Contact dynamics between different MLO pairs were analyzed using custom Python scripts based on time-resolved pairwise contact data tables acquired from MLO tracking and contact detection step above. Two complementary metrics, total contact dwell time and contact event counts per pair of MLOs, were summarized from the data tables across all 10 MLO pairwise contact combinations. Total contact dwell time was defined as the cumulative duration of all contact events for a given MLO pair overserved during imaging period. Contact event count was defined as the number of discrete contact events observed over the same period and was used as a measure of contact frequency.

MLO contact dynamics was first assessed qualitatively using line plot representations derived directly from contact data tables. For each MLO pair, contact states were mapped across the entire imaging duration, where continuous contact between two MLOs was represented by a solid line and interruptions in contact were represented by gaps. Each line therefore reflects the full temporal profile of an individual interaction event between a MLO pair, demonstrating both the duration and continuity of contact over time.

Additionally, as a more quantitative measure, contact dynamics were further characterized using two-dimensional kernel density estimation (2D-KDE) analysis^58^, resulting in a characteristic probability density map representing the unique interaction dynamics of each MLO pair. Contact events counts (X axis) and total contact dwell time (Y axis) were jointly used to construct a 2D distribution of interaction events in which each MLO pair was represented as a single data point within this 2D space. The collection of these data points was then used to capture the characteristic contact dynamics of each pairwise MLO combination.

Gaussian kernels were applied to the datasets of each pairwise MLO combination to estimate continuous probability density distributions without additional binning. Smoothing of the density estimate was controlled by the kernel bandwidth, which was determined automatically using Scott’s rule based on sample sizes^59^. Density values were evaluated across the full range (contact events counts: 0-12; total contact dwell time: 0-600) and normalized to the maximum value within each pairwise MLO combination, resulting in a relative density map ranging from 0 to 1. The 2D-KDE results were then visualized as contour lines (5%, 25%, 50%, and 75% of the maximum density) and filled contour regions projected onto the XY plane. For better visualization, a third dimension (Z axis) was introduced. This additional Z axis represents the fraction of total contact events corresponding to specific interaction states for a given pairwise MLO combination. These values were visualized as columns positioned over the corresponding regions in the XY plane.

Comparing across different MLO pairs, MLO pairs occupying regions of low contact event number (≤ 4) but long dwell times (≥ 200 s) were classified as stable interactions. Interactions exhibiting > 4 contact events were classified as transient interactions, reflecting repeated contact formation and separation events. Interactions exhibiting ≤ 4 contact events and < 200 s cumulative dwell time were classified as no/stochastic interactions.

#### MLO nearest-neighbor surface-to-surface distances

Nearest-neighbor surface-to-surface distance analysis between MLOs was performed using custom Python scripts on segmented binary mask datasets. First, individual-cell Airyscan images were segmented to yield composite mask dataset (4D, Z, C, Y, X) according to method described above. Subsequently, for each MLO objects identified from a given channel (e.g., CB in green channel), the minimum surface-to-surface distance to all objects in the target channel were calculated. For each source MLO, the surface was approximated as the outer one voxel shell obtained by subtracting a one-step 3D binary erosion from the full object mask. The nearest-neighbor distance to a target MLO was defined as the minimum Euclidean distance from the surface of the source MLO to any voxel in the target MLO mask. Distances were computed using a 3D Euclidean distance transform while accounting for the different pixel sizes in the XY and Z dimensions of the image (XY = 35 nm per pixel; Z = 130 nm per slice). Thus, the reported value represents the shortest surface-to-surface distance from each source MLO to the nearest object in the target channel. The shortest distance value between source and target MLO was recorded as the nearest-neighbor surface-to-surface distance, with overlapping or directly contacting objects assigned a distance of 0.

#### Nearest-neighbor MLO probability analysis

To characterize the spatial relationships among nuclear MLOs, nearest-neighbor surface-to-surface distance measurements obtained as described above were further analyzed. For each reference MLO object, the nearest neighboring MLO of a different type was identified, and the corresponding nearest-neighbor distance was recorded. Distance measurements were subsequently pooled across all analyzed cells for each pairwise MLO combination.

Cumulative probability plots were generated by calculating the fraction of neighboring MLOs located within a distance r from a reference MLO. These plots provide a quantitative measure of the likelihood that two MLO types are found in close spatial proximity, with steeper curves indicating stronger spatial association. To visualize the distribution of nearest-neighbor distances, measurements were grouped into 0.5 µm intervals and converted into probability density distributions. Distance distributions were smoothed using Gaussian kernel density estimation (bandwidth = 0.5) to generate continuous probability density curves.

#### Spatial distribution analysis of MLOs

The spatial organization of nuclear MLOs was quantified using Imaris. Individual MLOs were first reconstructed as 3D surfaces from Airyscan image stacks as described above. For each cell, the geometric center of the nuclear speckle was used as the reference center for radial distribution analysis within the nucleus. The radial intensity distribution of each MLO relative to this reference center was subsequently quantified. For axial distribution analysis, the Z-slice corresponding to the maximum intensity of the nuclear speckle was defined as the axial center of the nucleus. The intensity distribution of each MLO along the Z-axis relative to this reference plane was quantified. For population-level analyses, measurements from all objects of the same MLO type within each cell were pooled to obtain per-cell averages. Cell-to-cell variability was calculated as the coefficient of variation (CV = standard deviation / mean) across individual cells.

#### MLO trajectory correlation analysis following Actuator perturbation

To quantify coordinated movement between MLOs during nuclear deformation induced by Actuator-mediated nuclear actin assembly, centroid positions of segmented HLB, CB, and PML-NB masks were determined at each time point from time-lapse image sequences. For each MLO, displacement vectors between consecutive frames were calculated from centroid positions using a custom Python script. Displacement correlation was quantified using Pearson correlation analysis of stepwise displacement magnitudes between HLB & CB and between PML-NB & HLB or PML-NB & CB pairs. Directional correlation was quantified using cosine similarity of stepwise displacement vectors. High Pearson correlation values together with high cosine similarity values indicate synchronized MLO movements in both displacement magnitude and direction, consistent with coordinated movement expected from stable MLO contacts during nuclear deformation, whereas lower values indicate reduced coordination between MLO trajectories.

#### SPT analysis

Single-particle tracking (SPT) analysis was initially performed using Nikon NIS-Elements software. SmD1-Halo or NLS-Halo single particles were detected using the built-in spot detection function, with parameters set to spot typical diameters of 0.50 µm and a contrast threshold of 14.5. Spot detection was applied across all frames of the SPT dataset. Subsequently, detected spots were linked into trajectories using the object tracking function. Tracking was performed using a random motion model, with maximum 2 frames gap allowed. Additional parameters included a standard deviation multiplication factor of 2, and a maximum object speed of 200 µm/s. The resulting trajectory data, along with the calculated mean squared displacement (MSD), were exported for downstream quantitative analysis.

To classify particle trajectories relative to nuclear substructures, CB and NS masks were first generated from pre-SPT reference images. CB objects were subsequently categorized into two groups based on their spatial relationship with NS masks: (1) NS-contact CBs, defined as CB masks directly overlapping or contacting NS masks, and (2) NS-separate CBs, defined as CB masks without detectable contact with NS masks. SPT tracks whose coordinates overlapped with CB masks were retained and further separated into two populations: tracks associated with NS-contact CBs and tracks associated with NS-separate CBs. To ensure robust trajectory analysis, only tracks with a minimum duration of greater than 5 frames were retained for further analysis. For each trajectory, the effective diffusion coefficient (*D*_eff_) was calculated by linear fitting (MSD = a + bt) of the first four MSD points against time plot, where *D*_eff_ = b/4 for two-dimensional diffusion. The anomalous coefficient (α) was obtained from log_10_-log_10_ fitting of MSD versus time.

All *D*_eff_ and α values from the NS-contact-CB and NS-separate-CB groups were then pooled for population-level analysis similar to work from Ding *et al.*^60^ The pooled dataset was first visualized by plotting *D*_eff_ versus α, in which individual trajectories were displayed as scatter points and the overall distribution was represented by a two-dimensional kernel density estimate (KDE) contour map. Smoothing of the density estimate was controlled by a Gaussian kernel with bandwidth determined using Scott’s rule. Contour lines were drawn at 10%, 35%, 55%, 75%, and 95% of the maximum estimated density. The distributions of *D*_eff_ and α were further analyzed fitting a two-component Gaussian mixture model (GMM) in log_10_ and linear scale, respectively.

#### Characterization of MLO volume, surface area and moving rate

Quantitative characterization of MLO volume, surface area, and dynamic movement was performed using the Imaris software suite. Raw TIFF images (5D stacks, T, Z, C, Y, X) were imported into Imaris software for surface rendering and analysis. The Surface module was used to reconstruct 3D MLO surfaces from raw TIFF images using Otsu’s thresholding values as cutoff, allowing precise measurement of individual object volumes and surface areas. To assess dynamic behavior, time-lapse 3D TIFF dataset was analyzed using the surface tracking module in Imaris, which automatically detected and linked objects across sequential time points using a nearest-neighbor algorithm. The moving rate of each MLO was calculated based on changes in MLO object centroid positions across consecutive time points that are 20 seconds apart.

#### Observed and random pairwise MLO association probability

Pairwise MLO contact frequency was calculated as the number of observed contacts divided by the total possible contacts for a given MLO pair within each cell. For most MLO combinations, the total possible contacts were defined as the smaller number of objects present between the two MLO types. Because NS was segmented as a single object per cell, NS-associated contact frequencies were normalized to the number of the opposing MLO type. For nucleolus & PML-NB interactions, contact frequencies were normalized to the number of PML-NBs because a single nucleolus can simultaneously contact multiple PML-NBs. Contact enrichment or depletion was determined by comparing observed contact frequencies to randomized contact frequencies generated from spatial randomization analysis described below.

MLO randomization was performed to estimate pairwise MLO association probabilities using a custom Python script. Binary 3D masks representing segmented MLOs from real 3D images were first imported into the analysis. Individual MLO objects were randomly repositioned within the nuclear mask while preserving their original shapes and sizes, with placements accepted only if fully contained within the nuclear volume. For each dataset, 100 independent randomizations were performed, and randomized source objects were compared against fixed target masks using the same contact-detection criteria described above. Random association probability for each pairwise MLO combination was calculated as the total number of randomized contacts divided by the total number of successfully placed source objects across all iterations. Results were summarized as mean randomized contact counts and association probabilities for each pairwise MLO combination.

#### Interactome similarity analysis

To quantify the cell-to-cell similarity of MLO interaction patterns and interaction dynamics, pairwise MLO contact measurements were represented as vectors and compared using cosine similarity analysis. For MLO interaction pattern analysis, each cell was represented as a 10-dimensional vector containing the contact counts for all ten pairwise MLO combinations. Cosine similarity was subsequently calculated between all possible cell pairs to generate a cell-by-cell similarity matrix.

For MLO contact dynamics similarity analysis, two 10-dimensional vectors were generated for each cell, corresponding to the 10 possible pairwise MLO combinations: a contact-count vector and an average dwell-time vector. The contact-count vector contains the number of unique MLO contact pairs detected for each MLO combination during the 10-minute imaging period. For MLO combinations in which no contacts were observed, a value of zero was assigned. The average dwell-time vector contains the mean duration of individual contact events for each MLO combination. Cosine similarity was subsequently calculated between all contact-count or average dwell-time vectors to quantify cell-to-cell similarity in MLO contact dynamics and visualized as heatmaps. Specifically, for average dwell-time analysis, pairwise comparisons between cells were performed only using MLO combinations in which contacts were detected in both cells.

In both cell-to-cell similarity analysis for interaction patterns and interaction dynamics, cosine similarity values range from 0 to 1, where larger values indicate higher cell-to-cell similarities.

### 6. Immunoblot

Cells were lysed in cold RIPA buffer (89900, ThermoFisher) supplemented with cOmplete^TM^ Mini Protease Inhibitor Cocktail (11836170001, Sigma Aldrich). Cell lysate was cleared out by centrifugation at 21,000g for 15 min at 4 °C. Protein concentrations were determined using the Pierce^TM^ 660nm Protein Assay Kit (22662, ThermoFisher), and equal amounts of total protein were mixed with 5x SDS-PAGE protein loading buffer and boiled at 95°C for 5 minutes. Gel electrophoresis was performed using Mini-PROTEAN electrophoresis system (Bio-Rad) on a 4-12% gradient gel (M00654, GenScript). Proteins were then transferred to Odyssey Nitrocellulose Membrane (926-31090, LI-COR Biosciences) at 100 V for 90 min using ice-cold transfer buffer (25 mM Tris, 192 mM glycine, pH 8.3, 20% methanol v/v%). Membranes were blocked in 5% non-fat milk in TBST (20 mM Tris-HCl, 150 mM NaCl, 0.1% Tween-20) for 1 hour at room temperature, and incubated overnight at 4°C with primary antibodies diluted in blocking buffer. After repeated washing, membranes were incubated with fluorophore-conjugated secondary antibodies in blocking buffer for 1 hour at room temperature, and subsequently imaged using Odyssey M (LI-COR Biosciences). The antibodies used were anti-Coilin antibody [IH10] (ab87913, 1:5000 dilution), anti-PML antibody [EPR16792] (ab179466, abcam, 1:5000 dilution), anti-SRRM2 polyclonal antibody (PA5-66827, ThermoFisher, 1:2000 dilution), anti-CASP8AP2 polyclonal antibody (HPA053573, Millipore Sigma, 1:10000 dilution), anti-NPM1 monoclonal antibody (32-5200, Invitrogen, 1:1000 dilution), anti-β-actin antibodies (A1978, Millipore Sigma, 1:10000 dilution; 8457, Cell Signaling, 1:10000 dilution), anti-Vinculin monoclonal antibody (V9131, Millipore Sigma, 1:5000 dilution), IRdye 800CW donkey anti-mouse (926-32212, LI-COR Biosciences, 1:10000 dilution), IRdye 680RD goat anti-mouse (926-68070, LI-COR Biosciences, 1:10000 dilution), IRdye 800CW goat anti-rabbit (926-32211, LI-COR Biosciences, 1:10000 dilution), and IRdye 680RD goat anti-rabbit (926-68070, LI-COR Biosciences, 1:10000 dilution).

### 7. Immunofluorescence

Cells were seeded on #1.5 round glass coverslips in 24 well plate and fixed in 4% formaldehyde (12606S, Cell Signaling Technology) in PBS for 10 min. Following fixation, cells were permeabilized using 0.1% Triton X-100 in PBS for 15 minutes, and blocked with 3% bovine serum albumin (BSA) in PBS for 1 hour at room temperature. Primary antibodies were diluted in 1% BSA in PBS and added to coverslips for overnight incubation at 4°C. On the following day, coverslips were washed three times with PBS and incubated with Alexa Fluor-conjugated secondary antibodies (Invitrogen) diluted in 1% BSA in PBS for 1 hour at room temperature, protected from light. After additional washes, DNA was counterstained with Hoechst 33342 (62249, ThermoFisher) and mounted onto glass slides with Vectashield antifade mounting medium (H-1000-10, Vector Laboratories).

### 8. Drug treatment assays

#### 1,6-hexanediol treatment

HeLa cells and “*Rainbow Nucleus*” cells were seeded on 35 mm glass-bottom dishes (D35-20-1.5N, Cellvis) and cultured under standard conditions. 2 μg of LSS-mKate 2-NPM1 plasmid was transfected into the “*Rainbow Nucleus*” cells for 14 hours prior to the drug treatment. 100 nM Janelia Fluor 646 HaloTag ligand (a gift from Luke Lavis Lab) was added to the culture medium 2 hours prior to the drug treatment. Prior to the drug treatment, the culture medium was replaced with FluoroBrite™ DMEM (A1896701, Gibco) supplemented with 10% fetal bovine serum (26140079, Gibco), 1% GlutaMAX™ (35050061, Gibco), and 1% Penicillin-Streptomycin (15140122, Gibco). Epifluorescence imaging was conducted using a Nikon Ti2 Eclipse microscope equipped with a 100 x/ 1.49 NA oil-immersion objective, with cells maintained under 37°C with 5% CO2 in a stage-top incubator (Tokai-Hit). MLOs were excited by a D-LEDI light source. Images were acquired using the following excitation and filter settings: PML-NB, 385 nm (3% power, 50 ms exposure) with a DAPI filter; CB, 475 nm (3% power, 100 ms exposure) with a FITC filter; nucleolus, 475 nm (3% power, 200 ms exposure) with a TRITC filter; NS, 550 nm (2% power, 100 ms exposure) with a TRITC filter; and HLB, 621 nm (5% power, 100 ms exposure) with a Cy5 filter. These excitation conditions (LED power and exposure times) were verified to produce minimal photobleaching, ensuring that fluorescence changes observed during treatment were not attributable to imaging-induced signal loss. For perturbation experiments, 1,6-hexanediol (88571, Sigma-Aldrich) was added to a final concentration of 1% (v/v), and time-lapse imaging was initiated immediately at a rate of 1 frame per minute for a total duration of 60 minutes.

#### RNA polymerase inhibitor treatment

“*Rainbow Nucleus*” cells were prepared under the same conditions as described above for 1,6-hexanediol treatment. CX-5461 (Pol I inhibitor, HY-13323, MedChemExpress), triptolide (Pol II inhibitor, HY-32735), and ML-60218 (Pol III inhibitor, HY-122122, MedChemExpress) were added to the culture medium for 3 hours. Final concentrations of Pol inhibitors in medium were 0.5 μM for CX-5461, 0.5 μM for triptolide, and 40 μM for ML-60218, respectively. After treatment, cells were fixed with 4% formaldehyde (12606S, Cell Signaling Technology) in PBS for 10 min prior to imaging. Multi-spectral Airyscan imaging was performed using a Zeiss LSM900 confocal microscope, with acquisition parameters identical to those described in the *Image Acquisition* section.

#### 5-ethynyl uridine (5-EU) incorporation assay

To assess the effects of RNA polymerase inhibition on nascent RNA synthesis, we performed 5-EU incorporation assays. Cells were first treated with CX-5461 (0.5 µM), triptolide (0.5 µM), or ML-60218 (40 µM) for 3 hours. Following the 3-hour inhibitor treatment, 1 mM of 5-EU was added directly to the culture medium for additional 30 minutes to be incorporated into the newly synthesized RNA. Cells were subsequently fixed with 4% formaldehyde and processed using the Click-iT™ RNA Alexa Fluor™ 594 Imaging Kit (C10330, Thermo Fisher Scientific) following the manufacturer’s protocol. Following labeling, cells were imaged using a Zeiss LSM900 confocal microscope. Mean 5-EU fluorescence intensities were measured within the nucleolus and surrounding nucleoplasm, and the nucleolus-to-nucleoplasm intensity ratio was calculated for each cell as a quantitative measure of relative nascent RNA synthesis within the nucleolar compartment.

### 9. RNA immunoprecipitation qPCR (RIP-qPCR)

HeLa Coilin-mStayGold KI, mScarlet3-SRRM2 KI cells were seeded in tissue culture dishes 1 day before transfection and reached ∼70% confluency at the time of transfection. Cells were transfected with 3.0 μg SNRPD1-HaloTag or NLS-Halo control plasmid together with 9.0 μg empty pUC57-Mini vector (GenScript) using Lipofectamine 3000 Transfection Reagent (L3000015, Thermo Fisher Scientific) for 16 hours. Cells were then harvested at ∼10L cells per condition. After washing with ice-cold DPBS, cells were lysed in IP lysis buffer containing 20 mM HEPES, pH 7.9, 250 mM KCl, 2 mM MgCl_2_, 0.25% NP-40, 10% glycerol, 1 mM DTT, with freshly supplemented with EDTA-free protease inhibitor (cOmplete^TM^ Mini Protease Inhibitor Cocktail, 11836170001, Sigma Aldrich) and RNase inhibitor (SUPERase·In™ RNase Inhibitor, AM2694, Thermo Fisher Scientific). Cell lysates were incubated on ice for 15 min and passed through a 26G needle 10 times, followed by centrifugation at 21,000 × g for 15 min at 4 °C. A 50 μL aliquot of the supernatant was reserved as input, and the remaining supernatant was incubated with HaloTag antibody (anti-HaloTag pAb, G9281, Promega, 2 µg per 100 µL beads)-conjugated Protein G magnetic beads (Dynabeads™ Protein G, 10004D, Thermo Fisher Scientific) for 4 hours at 4°C with gentle shaking. Beads were washed 3 times with cold wash buffer containing 20 mM HEPES, pH 7.9, 150 mM KCl, 2 mM MgCl₂, 0.02% Tween-20, and 10% glycerol, freshly supplemented EDTA-free protease inhibitor and RNase inhibitor. RNA from both input and immunoprecipitated samples was extracted in TRIzol^TM^ reagent (15596026, Thermo Fisher Scientific). The subsequently RNA purification was conducted according to standard manufacturer’s instructions.

Following RNA purification, cDNA was synthesized using the High-Capacity RNA-to-cDNA Kit (4388950, Thermo Fisher Scientific) according to the manufacturer’s instructions. qPCR was carried out on a QuantStudio 3 Real-Time PCR System using PowerUp SYBR Green Master Mix (A25742, Applied Biosystems) with standard cycling and melt curve settings. qPCR signals for each target snRNA were normalized to 7SK RNA within the same sample, and relative enrichment was determined by comparison to the NLS-Halo control condition.

### 10. Statistical analysis

Statistical analysis was performed using GraphPad Prism software. Significance levels were reported using standard GraphPad notation: ns (not significant, p > 0.05), * (p ≤ 0.05), ** (p ≤ 0.01), *** (p ≤ 0.001), and **** (p ≤ 0.0001).

### 11. Gene fragments

#### Coilin donor repair fragments

caggctggtcttgaactcctgactttagatgatccgactgccttggcctcccagagtgctaggattataggcgtgagctaccatgcctggcctctttttttttgttttgttttttaattaaaaaaaataatagagatggggtctcactaggtttcccaggctgatcttgaactcctgggctcaagcatttctccccctcggccttccaaattgctgggattacaggtgtgagccacagcgcctggcctaaagagttactatactgcaattgaactcagtttattggaacccaaattattcccgttcttcaatcattctacacagacggaaaatttacgtggctgaattaaccggtatgacctc aaaatgaagtgcctaaactgggtattttgtctttttcctcagatcactgtattttggaaagagttgattgatcc<u>aagactgattattgaatctc</u>c aagtaacacatcaagtacagaacctgccggcggtggcggatcgatcgaattcctgcagcccgggggatccatggtgtctacaggcgaggagctgtttaccggcgtggtgcccttcaagttccagctgaagggcaccatcaacggcaagagcttcaccgtggaaggcgagggcgagggcaatagccacgagggcagccacaaaggcaagtacgtgtgcaccagcggcaaactgccaatgtcttgggccgccctgggaactagcttcggctatggcatgaagtactacaccaagtaccccagcggcctgaagaactggttccacgaggtgatgcccgagggctt cacctacgacagacacatccagtacaagggcgacggcagcatccacgccaagcaccagcacttcatgaagaacggcacctaccacaacatcgtggagttcaccggccaggacttcaaggagaacagccccgtgctgaccggcgacatggacgtgagcctgcccaacgaggtgcagcacatccccatagatgacggcgtggagtgcacagtgaccctgcagtaccctctgctgagcgacgagagcaagtgcgtggaggcctaccagaacaccatcatcaagcccctgcacaatcagccagcccccgatgtgccattccactggatcagaaagcagtacacccagagcaaggacgacaccgaggagagagaccacatcatccagagcgagaccctggaggcccacctgtaatgagtatgacctcttcaccttatagtttatgaatgtcttgtttgtgaaagtgactataacccaaactttttttattttttaaagaggatttggaagttgtatggatttttttgttatcttcactttactgcataggaaacaatctacctcatcatttaaaatgacatgggtgtcggttttgtagatctttggtttttttgtcaggtttaatttcagttaacaaaatgtaaaacatgacattccctgcagatattgttgtataccagtatggtttcttctctttctttaaatgtttttggccatcaagtagcagtcgtcagtaggagtttataataccaagaatgtgctgcgtatcttgtctcaataagttttaagtaacatttaaaaatattaaagcatgttatttgacctaattttttagcatttgagttgttccattaaatggagcatcttgtaaatttcaagtattttatacttgcaattgtt

5’ homology arm; linker sequence; mStaygold coding sequence; 3’ homology arm; <u>guide RNA targeting sequence</u>.

#### SRRM2 donor repair fragments

tgccacagatctgtgtctggtgtttgttgtgaccaggaggctttggttggttaccagaggttaggaattagagtagttggtaaacacaccacttgtgcattttggggccaaaggagaacaaaaagggacattgtggagagattttagaggaacaagatccttgtttgcctagggtggaaaaagaagaaaatttaggtgtagggagggattatttagaattctgtgcaagggcattgagagaaaggctgtaagaactgggctgtgatttggtagaaaggaaatttggaagggcaaagaagtggttttcttgggagaccaggaggatggtggagcattatacagcgaagacttagtgtgaagatcaatgttgaattagggatgaaggagagcagggacttgggtgtggggcggtaagtggttagtttggggccaggg gcctgacccgtgtctccccgctccccctcaggagcggtggtgccccccccgggcacg<u>gggccatg</u>gatagcaccgaggcagtgatc aaggagttcatgcggttcaaggtgcacatggagggctccatgaacggccacgagttcgagatcgagggcgagggcgagggccgcccctacgagggcacccagaccgccaagctgagggtgaccaagggtggccccctgcccttctcctgggacatcctgtcccctcagttcatgtacggctccagggccttcacgaagcaccccgccgacatccccgactactggaagcagtccttccccgagggcttcaagtgggagcgcgtgatgaacttcgaggacggcggcgccgtgtccgtggcccaggacacctccctggaggacggcaccctgatctacaaggtgaagctccgcggcaccaacttccctcctgacggccccgtaatgcagaagaagacaatgggctgggaagcatccaccgagcggttgtaccccgaggacgtcgtgctgaagggcgacattaagatggccctgcgcctgaaggacggcggccgctacctggcggacttcaagac cacctacagggccaagaagcccgtgcagatgcccggcgccttcaacatcgaccgcaagttggacatcacatcccacaacgagga ctacaccgtggtggaacagtacgaacgctccgtggcccgccactccaccggcggctccggtggctcctctagatcagaagga<u>taca acgggatc</u>gggctgccgacgccccggggcagcggcaccaacggctacgtccagcgcaacctgtccctggtgcggggccgccgg ggtgagcggcctgactacaagggagaggaggaactgcggcgcctggaggctgccctggtgaagcggcctaatcctgacatcctggaccacgagcgcaagcggcgcgtcgagctgcgatgcctcgagctggaggagatgatggaagagcaggggtgagggagagctgggggagagtcaagcactgaatgagtgcagagctgggggtgttaggtgggatgtatagggagcttagggtggttgaaagaggcctggcaaagagttgtggtaggggaggaggcagatgagctctagagaagtgaaaacagttaagggacagtgcagagtgggaaatgagaaggctgtgtgtctggggcatagatgggagcctgaggtactaaatggaggcacgtgggagaagggaggggccattgagg

5’ homology arm; mScarlet3 coding sequence; linker sequence; 3’ homology arm; <u>guide RNA targeting sequence</u>.

