## Supplementary Figures for "Rainbow Nucleus Charts Dynamic Interactome of Membrane-less Organelles"

### 1. Supplementary Figures

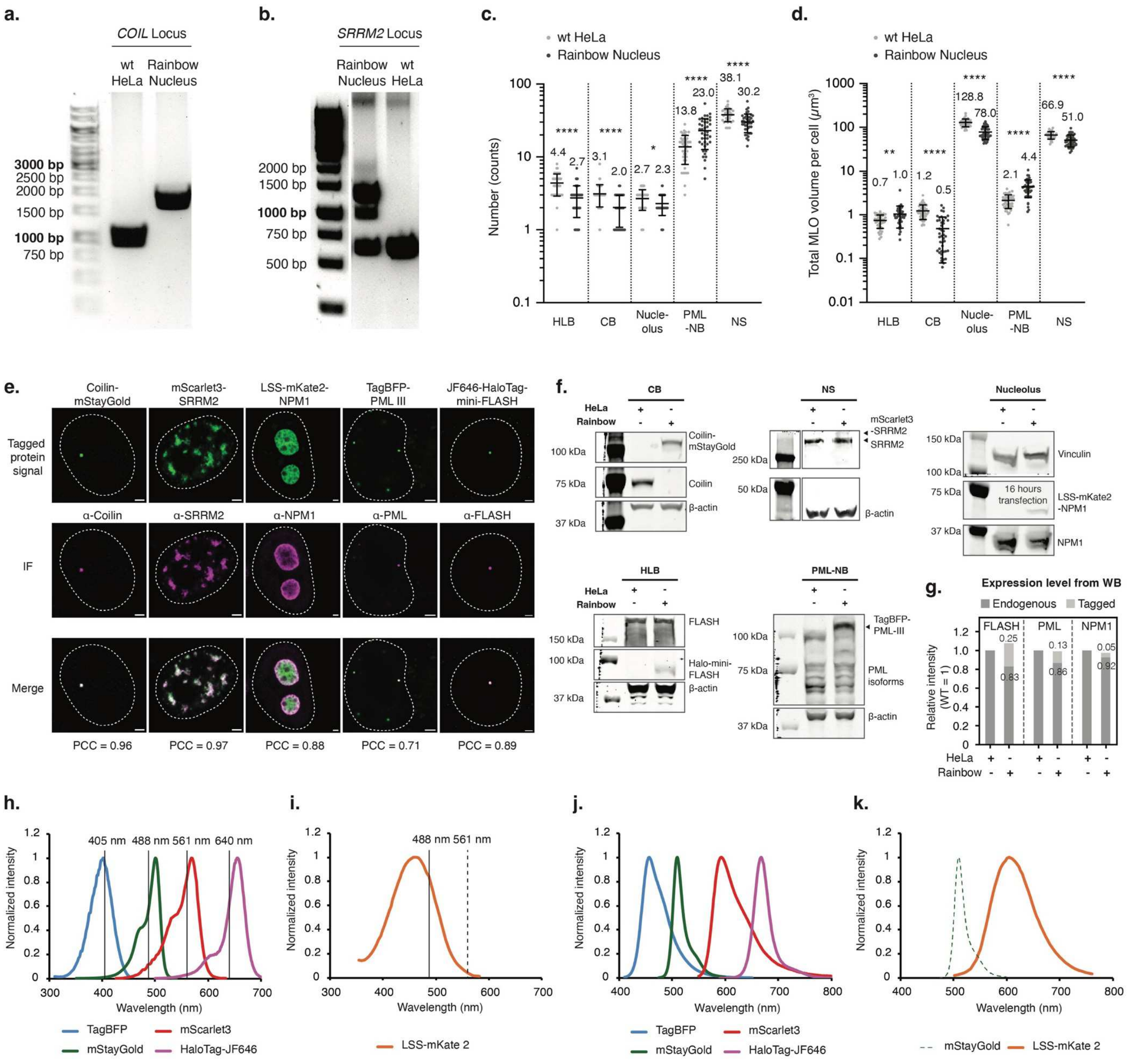

**Figure S1. Validation, characterization, and spectral imaging workflow of the “Rainbow Nucleus” cell line.**

**(a-b)** Genomic PCR gel images indicating the knock-in of mStayGold gene at the *COIL* locus (**a**), and the mScarlet3 gene at the *SRRM2* locus (**b**).

**(c-d)** Quantification of the number (**c**) and total MLO volume per cell (**d**) of individual nuclear MLOs in wild-type (wt) HeLa and “Rainbow Nucleus” cells. While statistical significances were observed, the overall changes in MLO abundance and volume remained relatively minor, indicating that introduction of the fluorescent labels did not substantially perturb MLO morphologies. Horizontal bars and labeled numbers indicate mean values; error bars indicate the standard deviations. Statistical comparisons were performed using unpaired two-tailed t tests. Significance levels were annotated as follows: $0.01 < p \leq 0.05$, *; $0.001 < p \leq 0.01$, **; $p \leq 0.0001$, ****. Sample sizes are as follows: HLB, n = 43 wt fixed HeLa cells; CB, n = 54 wt fixed HeLa cells; nucleolus, n = 45 wt HeLa cells; PML-NB, n = 43 wt HeLa cells; and NS, n = 35 wt HeLa cells. All comparisons were performed against n = 43 fixed "Rainbow Nucleus" cells.

**(e)** Validation of fluorescent reporters by immunofluorescence. Representative images showing fluorescence signals from tagged fluorescent proteins or HaloTag-JF 646 conjugate (top), immunofluorescence staining (middle), and merged images (bottom) for each MLO marker protein. Pearson's correlation coefficients (PCC) indicate colocalization between tagged reporters and immunofluorescence signal. Tagged protein signals are pseudocolored in green, and immunofluorescence signals in magenta for better contrast. Scale bars, 2 μm.

**(f)** Immunoblots showing the expression level of tagged reporters and corresponding endogenous proteins for each MLO marker in HeLa and in "Rainbow Nucleus" cells. β-actin or vinculin were used as loading controls.

**(g)** Relative expression levels of endogenous and tagged MLO marker proteins in "Rainbow Nucleus" compared with HeLa cells from (**f**).

**(h-i)** Reference excitation spectrum of TagBFP, mStayGold, mScarlet3, HaloTag-JF646 (**h**), and LSS-mKate2 (**i**). Solid vertical lines indicate laser lines used for imaging (405, 488, 561, and 640 nm), and the dashed line in (**i**) marks 561 nm for comparison of excitation efficiency relative to 488 nm for LSS-mKate2. Notably, 561 nm does not efficiently excite LSS-mKate2, thereby minimizing cross-excitation with mScarlet3.

**(j-k)** Reference emission spectrum of TagBFP, mStayGold, mScarlet3, HaloTag-JF646 (**j**), and LSS-mKate2 (**k**). The dashed emission spectrum in (**k**) indicates mStayGold emission for comparison with LSS-mKate2 spectrum. The spectral separation between mStayGold and LSS-mKate2 enables clear separation with minimal emission crosstalk.

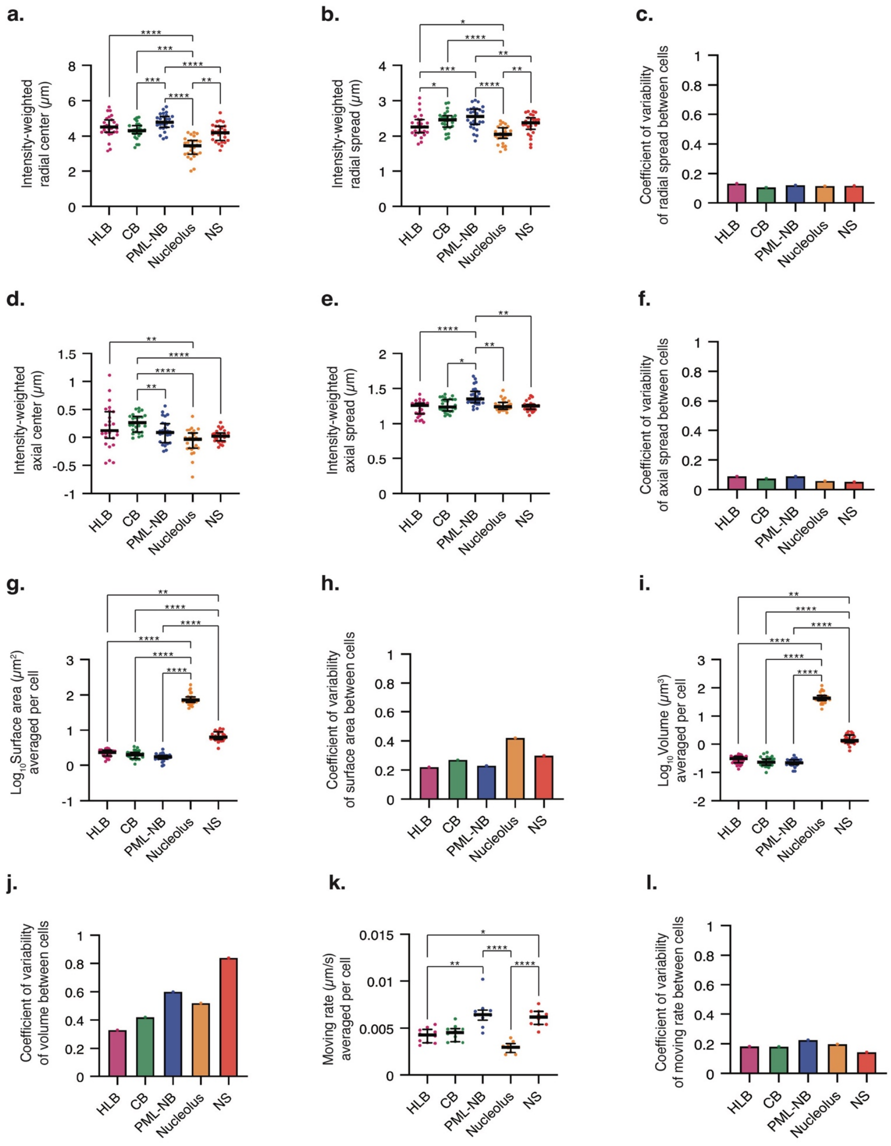

**Figure S2. Cell-to-cell variability in spatial organization, morphology, and dynamics of nuclear MLOs in "Rainbow Nucleus" cells.**

**(a-b)** Scatter plots showing the intensity-weighted radial center (**a**) and radial spread (**b**) of individual nuclear MLOs across cells. The radial center and spread were calculated relative to the nuclear center defined by the centroid of the NS signals in the radial dimension. Horizontal bars indicate medians; vertical whiskers indicate the interquartile range. Statistical comparisons were performed using Friedman tests followed by Dunn's multiple-comparisons tests. Only statistically significant comparisons are labeled; unlabeled comparisons indicate non-significant differences ($p > 0.05$). Significance levels were annotated as follows: $0.01 < p \leq 0.05$, *; $0.001 < p \leq 0.01$, **; $0.0001 < p \leq 0.001$, ***; $p \leq 0.0001$, ****. n = 27 fixed "Rainbow Nucleus" cells.

**(c)** Bar plots showing the coefficient of variability (CV) of radial spread across cells for each nuclear MLO from 27 fixed “Rainbow Nucleus” cells. CV values were calculated as the standard deviation divided by the mean of radial spread measurements across cells.

**(d-e)** Scatter plots showing the intensity-weighted axial center (**d**) and axial spread (**e**) of individual nuclear MLOs across cells. The axial center and spread were calculated relative to the nuclear center defined by the centroid of the NS signals in the axial dimension. Horizontal bars indicate medians; vertical whiskers indicate the interquartile range. Statistical comparisons were performed using Friedman tests followed by Dunn’s multiple-comparisons tests. Only statistically significant comparisons are labeled; unlabeled comparisons indicate non-significant differences ($p > 0.05$). Significance levels were annotated as follows: $0.01 < p \le 0.05$, *; $0.001 < p \le 0.01$, **; $p \le 0.0001$, ****. n values are the same as in (**a-b**).

**(f)** Bar plots showing the CV of axial spread across cells for each nuclear MLO from 27 fixed “Rainbow Nucleus” cells.

**(g-h)** Scatter plots showing per cell average surface areas (**g**) and corresponding CV of average surface areas across cells (**h**) for nuclear MLOs. Surface area measurements were derived from Imaris surface segmentation using Otsu thresholding. In (**g**), horizontal bars indicate medians; vertical whiskers indicate the interquartile range. Statistical significance was determined using Friedman tests followed by Dunn’s multiple-comparisons tests. Only statistically significant comparisons are labeled; unlabeled comparisons indicate non-significant differences ($p > 0.05$). Significance levels were annotated as follows: $0.001 < p \le 0.01$, **; $p \le 0.0001$, ****. CV values in (**h**) were calculated as the standard deviation divided by the mean surface area measurements. n values are the same as in (**a-b**).

**(i-j)** Scatter plots showing per cell average volumes (**i**) and corresponding CV of average volumes across cells (**j**) for nuclear MLOs. Volume measurements were derived from Imaris surface segmentation using Otsu thresholding. In (**i**), horizontal bars indicate medians; vertical whiskers indicate the interquartile range. Statistical significance was determined using Friedman tests followed by Dunn’s multiple-comparisons tests. Only statistically significant comparisons are labeled; unlabeled comparisons indicate non-significant differences ($p > 0.05$). Significance levels were annotated as follows: $0.001 < p \le 0.01$, **; $0.0001 < p \le 0.0001$, ****. CV values in (**j**) were calculated as the standard deviation divided by the mean volume measurements. n values are the same as in (**a-b**).

**(k-l)** Scatter plots showing the per cell average moving rates (**k**) and corresponding CV of average moving rates across cells (**l**) of nuclear MLOs across individual cells. Moving rates were calculated in Imaris using the surface speed function, which measures instantaneous speed based on changes in MLO object centroid positions across consecutive time points. In (**k**), horizontal bars indicate medians; vertical whiskers indicate the interquartile range. Statistical significance was determined using Friedman tests followed by Dunn’s multiple-comparisons tests. Only statistically significant comparisons are labeled; unlabeled comparisons indicate non-significant differences ($p > 0.05$). Significance levels were annotated as follows: $0.01 < p \le 0.05$, *; $0.001 < p \le 0.01$, **; $p \le 0.0001$, ****. CV values in (**l**) were calculated as the standard deviation divided by the mean moving rates. Moving rates in (**k-l**) were calculated in Imaris using surface speed function, which calculates instantaneous speed based on changes in MLO object centroid positions across consecutive time points, collected from 10 live “Rainbow Nucleus” cells across 31 time points (20 s interval; total duration 600 s).

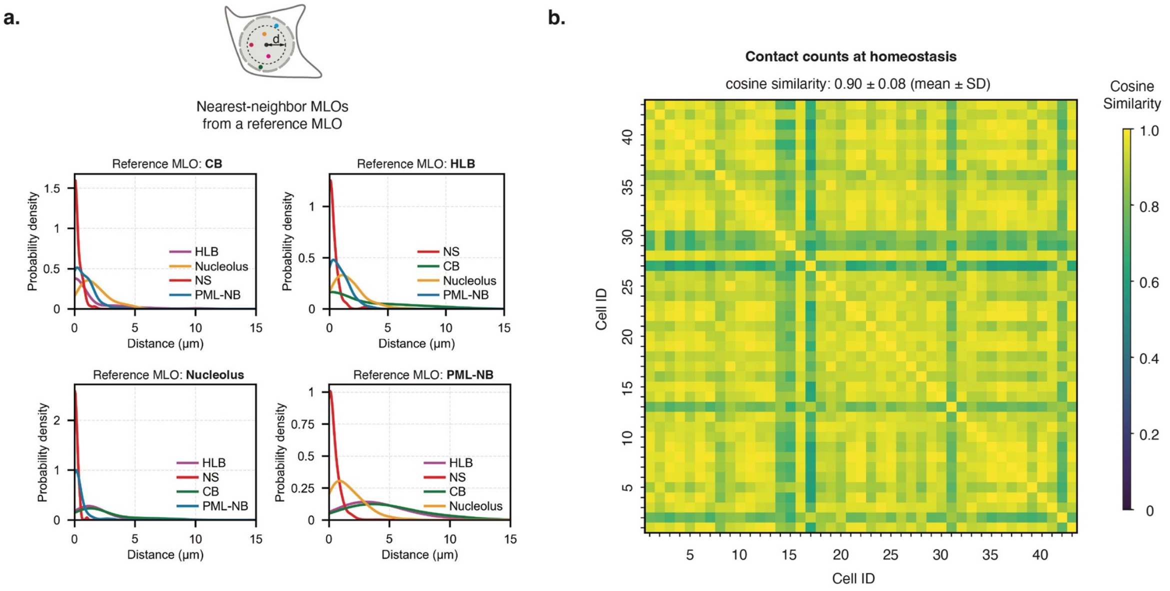


**Figure S3. Nuclear MLOs establish a structured interactome with high cell-to-cell consistency.**

**(a)** Probability density plots indicating the likelihood of reference MLO encountering the nearest MLO of a different type as a function of distance in “Rainbow Nucleus” cells. n = 118 for HLB, n = 87 for CB, n = 987 for PML-NB, and n = 98 for nucleolus, collected from 43 fixed “Rainbow Nucleus” cells. Histograms were constructed using a bin width of 0.5 μm and smoothed using kernel density estimation with a bandwidth of 0.5.

**(b)** Heatmap showing pairwise cosine similarity of pairwise MLO contact counts across 43 fixed “Rainbow Nucleus” cells. For each cell, contact counts were represented as a 10-dimensional vector spanning all 10 pairwise MLO interactions. Cosine similarity was calculated between corresponding vectors across all cells.

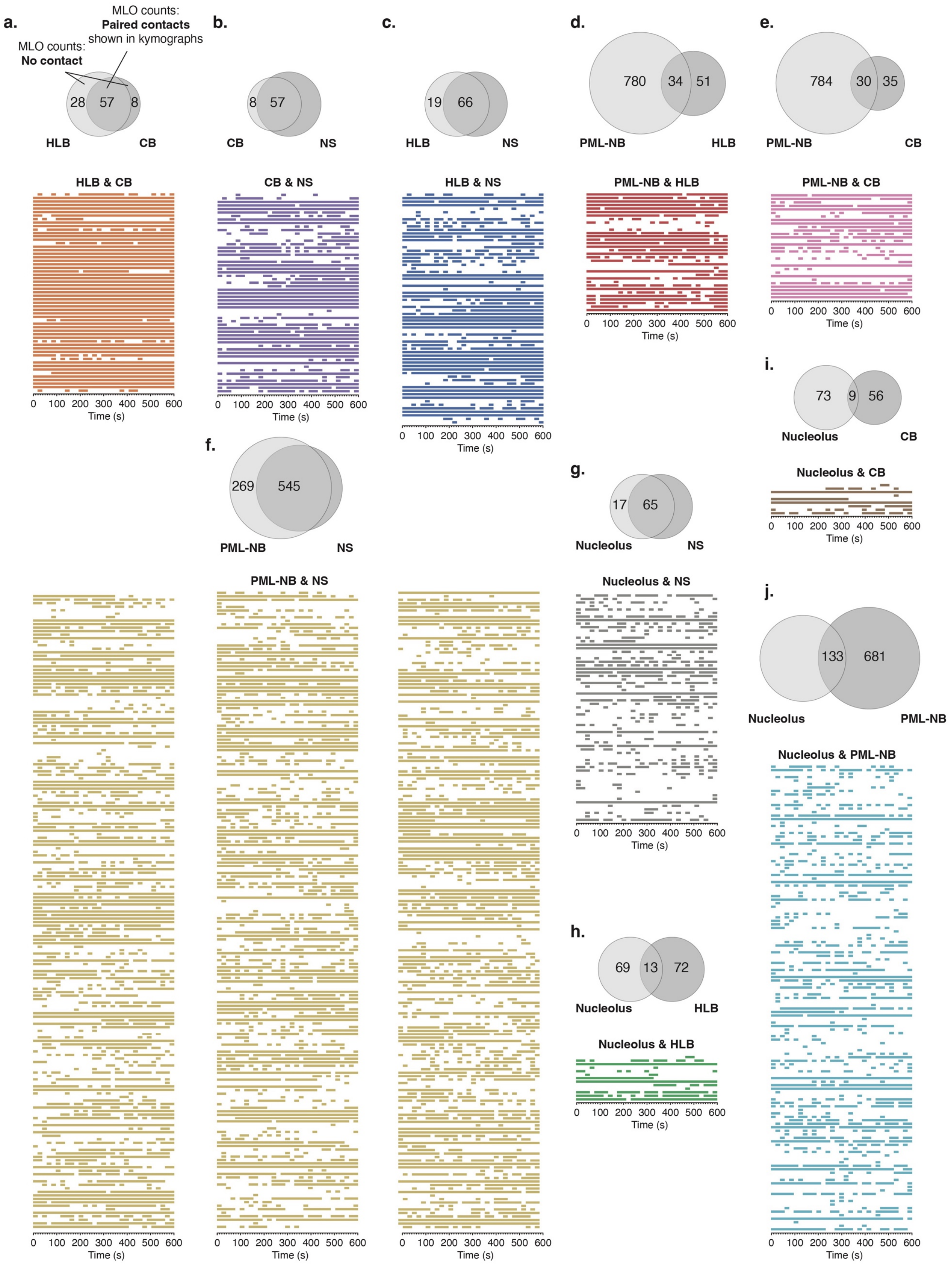

a.
MLO counts:
Paired contacts
shown in kymographs
MLO counts:
No contact
28
57
8
HLB
CB
HLB & CB
b.
8
57
CB
NS
CB & NS
c.
19
66
HLB
NS
HLB & NS
d.
780
34
51
PML-NB
HLB
PML-NB & HLB
e.
784
30
35
PML-NB
CB
PML-NB & CB
f.
269
545
PML-NB
NS
PML-NB & NS
g.
17
65
Nucleolus
NS
Nucleolus & NS
h.
69
13
72
Nucleolus
HLB
Nucleolus & HLB
i.
73
9
56
Nucleolus
CB
Nucleolus & CB
j.
133
681
Nucleolus
PML-NB
Nucleolus & PML-NB
0 100 200 300 400 500 600
Time (s)

**Figure S4. Quantification and kymograph visualization of pairwise nuclear MLO contact dynamics.**

Pairwise contact dynamics between (**a**) HLB & CB, (**b**) CB & NS, (**c**) HLB & NS, (**d**) PML-NB & HLB, (**e**) PML-NB & CB, (**f**) PML-NB & NS, (**g**) nucleolus & NS, (**h**) nucleolus & HLB, (**i**) nucleolus & CB, and (**j**) nucleolus & PML-NB in live “Rainbow Nucleus” cells. Top schematics summarize the number of MLOs without contact and those engaged in paired contacts for each MLO pair. Kymographs display contact events over time for individual MLO pairs, where each row represents a tracked MLO pair and continuous line segments indicate sustained contacts. These contact data were used for quantifications of probability density plots in **Fig. 4e-n**. Sample size, n = 32 live “Rainbow Nucleus” cells imaged over 10 minutes at 20-second intervals.

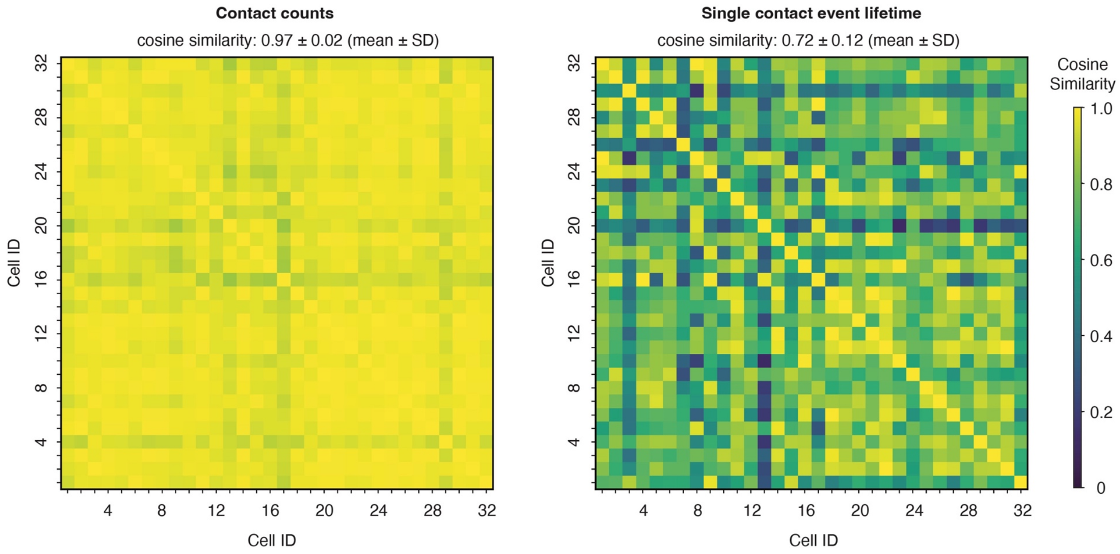


**Figure S5. Cell-to-cell variability analysis reveals conserved nuclear MLO contact dynamics.**

Heatmaps showing pairwise cosine similarity of pairwise MLO contact counts (left panel) and single contact event lifetime (right panel) across 32 live "Rainbow Nucleus" cells. For each cell, contact dynamics were represented as two 10-dimensional vectors spanning all 10 pairwise MLO interactions: a contact-count vector and a single-contact-event-lifetime vector. Cosine similarity was calculated between corresponding vectors across all cells. For single contact event lifetime similarity analysis, MLO pairs lacking detectable interactions in either cell were excluded from the pairwise comparison, and similarities were calculated only across shared observed interaction dimensions.

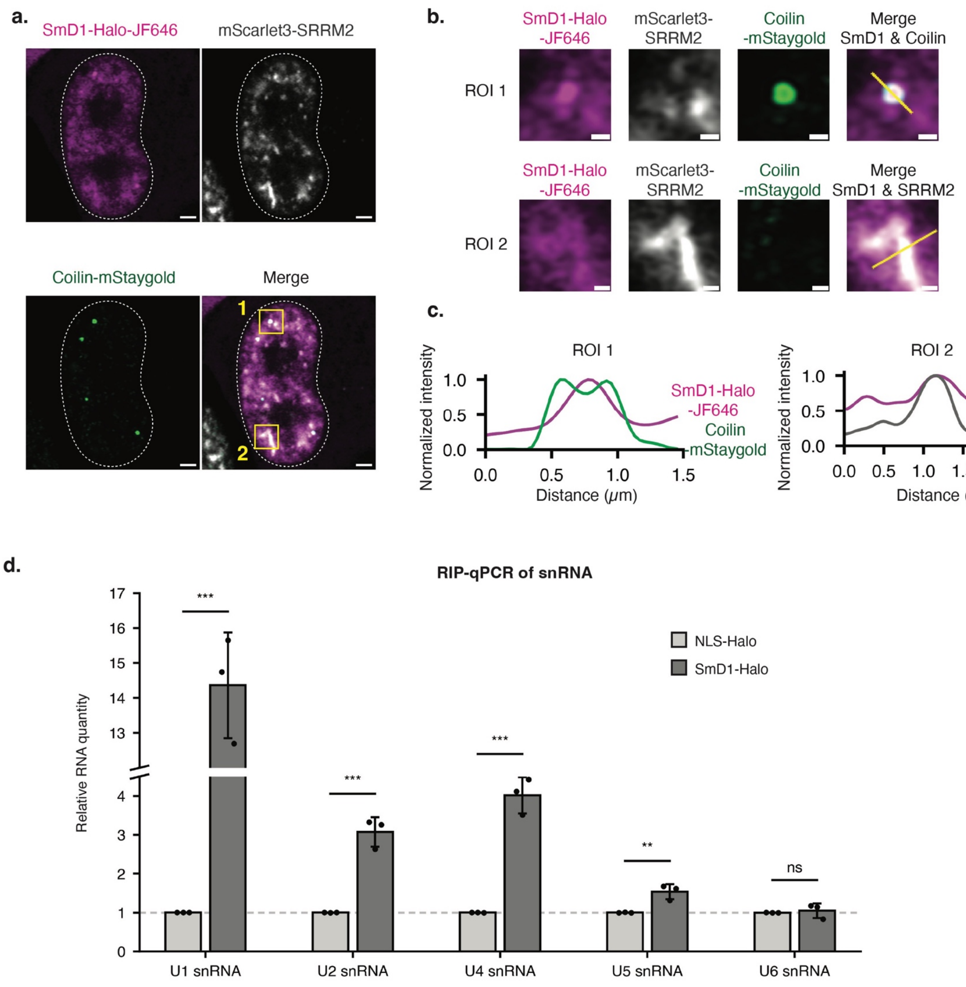


**Figure S6. SmD1-Halo protein localizes to CB and NS similarly to *bona fide* SmD1 and is incorporated into snRNP complexes.**

**(a-c)** Localization of SmD1-Halo relative to CBs and NSs. (**a**) Representative of a HeLa cell with endogenous mScarlet3-SRRM2 and Coilin-mStayGold knock-ins, together with transiently expressed SmD1-Halo, labeled with JF646, used for SmD1 localization analysis relative to CBs and NSs. Scale bars, 2 μm. (**b**) Magnified views of the boxed ROIs in (**a**), showing representative colocalization of SmD1-Halo with coilin signal (top) and SRRM2 signal (bottom). Scale bars, 0.5 μm. (**c**) Line profile analysis of normalized fluorescence intensity across the indicated regions in (**b**), showing spatial overlap of SmD1-Halo with coilin and SRRM2 signal.

**(d)** Bar graph showing snRNA enrichment following immunoprecipitation of Halo-tagged proteins from cells transiently expressing SmD1-Halo or NLS-Halo for 16 hours. snRNA associated with Halo-tagged proteins was isolated using anti-Halo immunoprecipitation, followed by qPCR quantification. Signals were normalized to 7SK RNA within each sample, and relative enrichment in SmD1-Halo was compared to the NLS-Halo control. Data are presented as mean ± SD. Statistical significance was assessed using two-tailed t-test. Significance levels were annotated as follows: $p > 0.05$, ns; $0.001 < p \leq 0.01$, **; $0.0001 < p \leq 0.001$, ***. Sample sizes, N = 3 repeats.

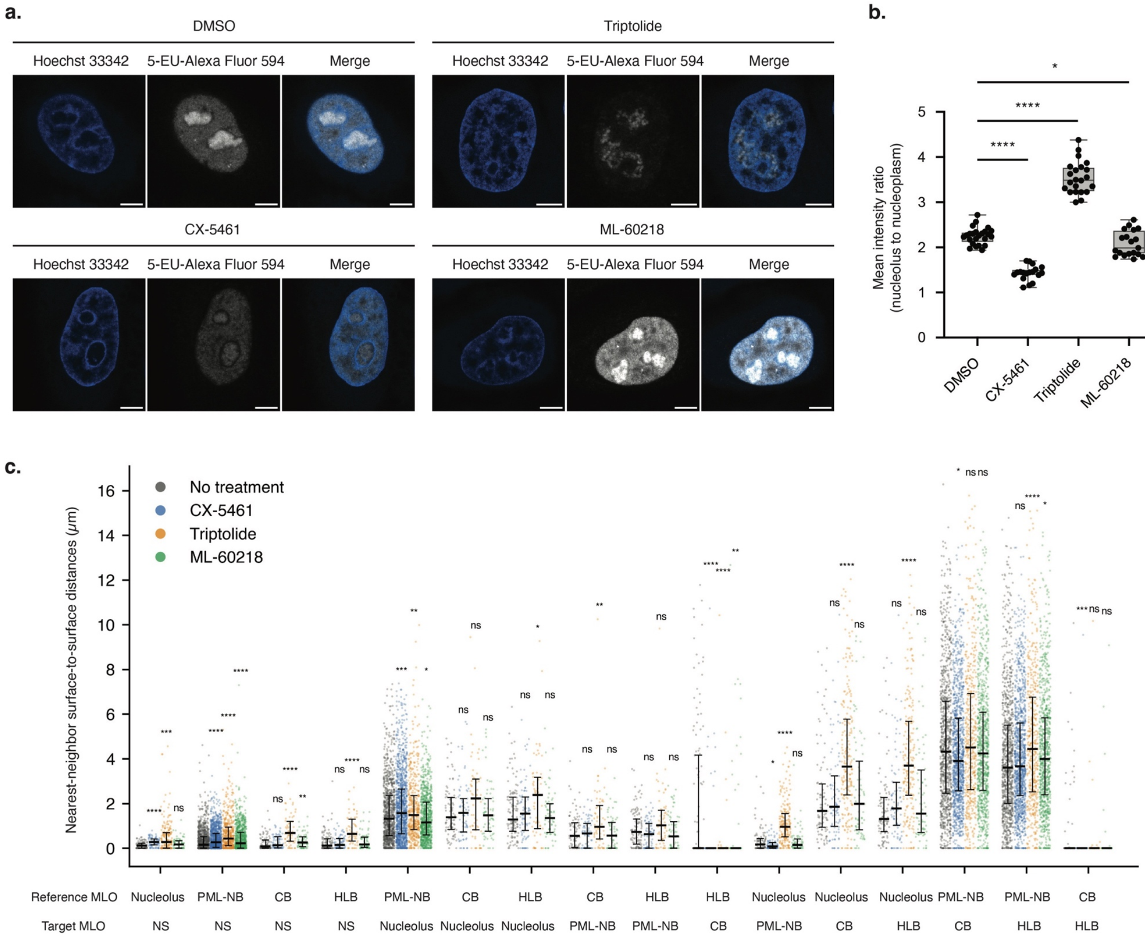


**Figure S7. Transcriptional perturbation reshapes nuclear MLO spatial organization.**

**(a)** Representative fixed cell images of 5-ethynyluridine (5-EU) incorporation in HeLa cells treated with DMSO, CX-5461 (0.5 μM), triptolide (0.5 μM), or ML-60218 (40 μM). Cells were treated with the indicated drugs for 3 hours, followed by 5-EU labeling for 30 min prior to fixation and AF594-azide labeling. Scale bars, 5 μm.

**(b)** Quantification of the mean intensity ratio of nucleolar to nucleoplasmic 5-EU signal in HeLa cells treated with DMSO, CX-5461 (0.5 μM), triptolide (0.5 μM), or ML-60218 (40 μM) for 3 hours. This ratio reflects the relative contribution of nucleolar (Pol I-dependent) versus nucleoplasmic (Pol II/III-dependent) transcription. Statistical significance was assessed using two-tailed t-test. Significance levels were annotated as follows: $0.01 < p \leq 0.05$, *; $p < 0.0001$, ****. Sample sizes, n = 24 fixed HeLa cells for DMSO treatment, n = 18 fixed Hela cells for CX-5461 treatment, n = 22 fixed HeLa cells for Triptolide treatment, and n = 19 fixed HeLa cells for ML-60218 treatment.

**(c)** Scatter plot showing the nearest-neighbor surface-to-surface distances between indicated nuclear MLO pairs in “Rainbow Nucleus” cells under no treatment, CX-5461 (0.5 μM), triptolide (0.5 μM), or ML-60218 (40 μM) conditions. Cells were treated with indicated drugs for 3 hours. Each point represents an individual distance measurement. Black horizontal lines indicate medians, and vertical whiskers indicate the interquartile range. Sample sizes are n = 118 (HLB), 87 (CB), 987 (PML-NB), and 98 (nucleolus), collected from 43 fixed “Rainbow Nucleus” cells for no-treatment condition; n = 82 (HLB), 75 (CB), 1198 (PML-NB), and 89 (nucleolus), collected from 33 fixed “Rainbow Nucleus” cells for CX-5461 treatment; n = 61 (HLB), 60 (CB), 495 (PML-NB), and 210 (nucleolus) from 30 fixed “Rainbow Nucleus” cells for triptolide treatment; and n = 70 (HLB), 63 (CB), 890 (PML-NB), and 74 (nucleolus) from 30 fixed “Rainbow Nucleus” cells for ML-60218 treatment. Statistical significance was assessed using two-tailed Mann-Whitney U tests comparing each transcriptional inhibition condition to no-treatment homeostasis controls. Significance levels were annotated as follows: $p > 0.05$, ns; $0.01 < p \leq 0.05$, *; $0.001 < p \leq 0.01$, **; $0.0001 < p \leq 0.001$, ***; $p \leq 0.0001$, ****.

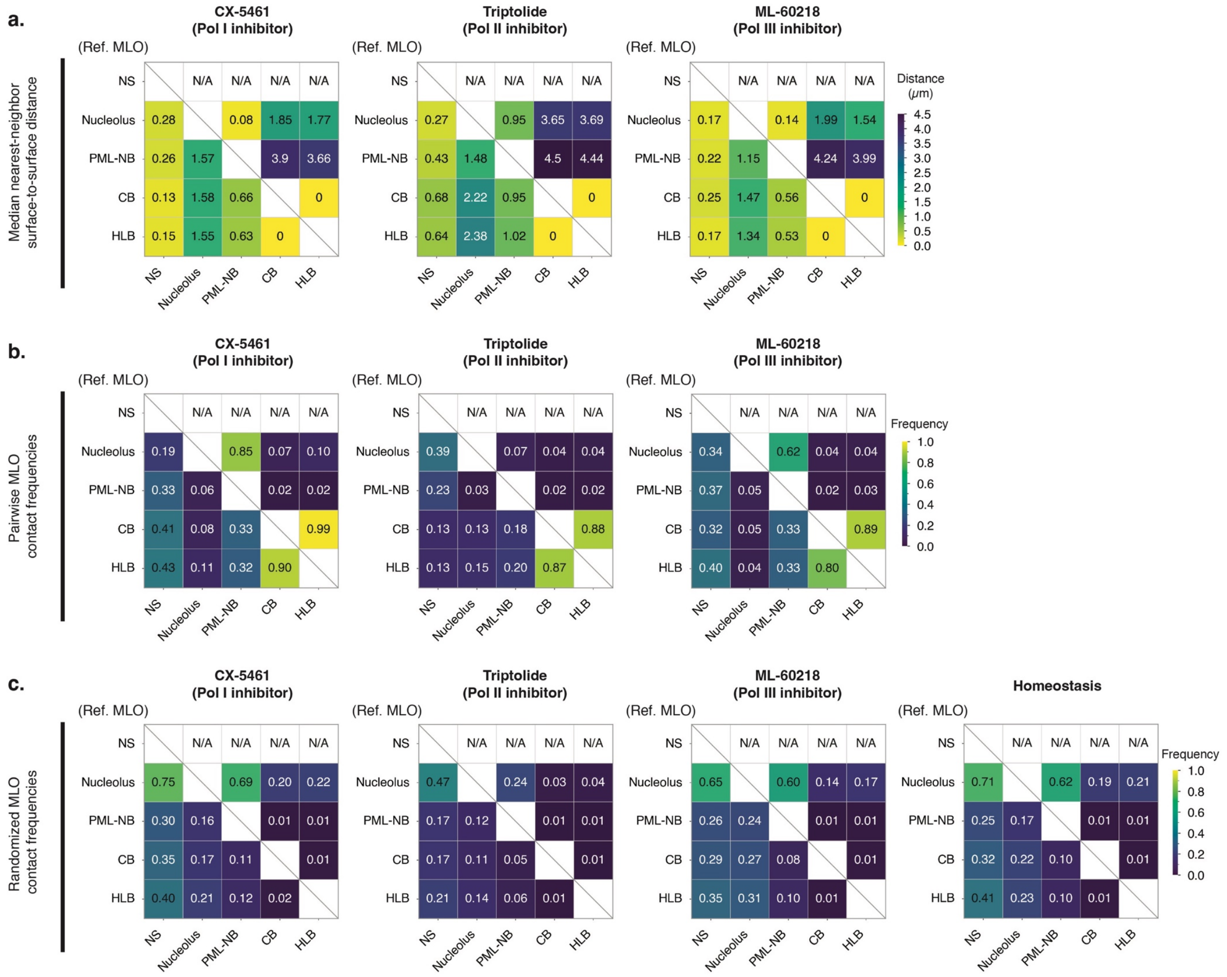


**Figure S8. Systematic mapping of pairwise nuclear MLO spatial relationships and contact frequencies under transcriptional inhibition.**

**(a)** Heatmaps of median nearest-neighbor surface-to-surface distances among nuclear MLOs in “Rainbow Nucleus” cells treated with CX-5461 (0.5 μM, left), triptolide (0.5 μM, middle), or ML-60218 (40 μM, right) for 3 hours. Color indicates the median distance between each reference MLO (row labels, left) and target MLO (column labels, bottom). n = 82 (HLB), 75 (CB), 1198 (PML-NB), and 89 (nucleolus), collected from 33 fixed “Rainbow Nucleus” cells for CX-5461 treatment; n = 61 (HLB), 60 (CB), 495 (PML-NB), and 210 (nucleolus) collected from 30 fixed “Rainbow Nucleus” cells for triptolide treatment; and n = 70 (HLB), 63 (CB), 890 (PML-NB), and 74 (nucleolus) collected from 30 fixed “Rainbow Nucleus” cells for ML-60218 treatment. NS were thresholded as a single, whole object per cell in the distance analysis.

**(b)** Heatmaps of pairwise MLO contact frequencies among nuclear MLOs in “Rainbow Nucleus” cells treated with CX-5461 (0.5 μM, left), triptolide (0.5 μM, middle), or ML-60218 (40 μM, right) for 3 hours. Color indicates the fraction of reference MLOs (row labels, left) in contact with target MLOs (column labels, bottom), relative to the total numbers of reference MLOs. Sample sizes are the same as in (**a**).

**(c)** Heatmaps of randomized pairwise MLO contact frequencies among nuclear MLOs in "Rainbow Nucleus" cells treated with CX-5461 (0.5 μM, left), triptolide (0.5 μM, middle), or ML-60218 (40 μM, right) for 3 hours. Color indicates the expected fraction of reference MLOs (row labels) in contact with target MLOs (column labels), calculated from repositioning MLO objects within the nuclear volume while preserving their shapes and sizes. For each treatment condition, randomization analysis was performed using segmented MLO masks derived from 10 drug-treated or homeostatic fixed "Rainbow Nucleus" cells.

### 2. Supplementary Tables

Table S1. List of oligonucleotides used in this work.

| | Target | Forward primer | Reverse Primer |
|---|---|---|---|
| qPCR primer | U1 snRNA | 5’-atacttacctggcaggggaga-3’ | 5’-gcagtcgagtttcccacatt-3’ |
| | U2 snRNA | 5’-atcgcttctcggccttttt-3’ | 5’-tattccatctccctgctcca-3’ |
| | U4 snRNA | 5’-tggcagtatcgtagccaatg-3’ | 5’-ctgtcaaaaattgccagtgc-3’ |
| | U5 snRNA | 5’-ctctggtttctcttcagatcgc-3’ | 5’-caaggcctcaaaaaattggg-3’ |
| | U6 snRNA | 5’-cgcttcggcagcacatatac-3’ | 5’-atggaacgcttcacgaattt-3’ |
| | 7SK RNA | 5’-catccccgatagaggaggac-3’ | 5’-caaatggaccttgagagcttg-3’ |
| sgRNA sequence | SRRM2 | 5’-gggccatgtacaacgggatc-3’ | |
| | Coilin | 5’-gagattcaataatcagtctt-3’ | |
| Genomic PCR | SRRM2 | 5’-gtgcaagggcattgagagaaaggc-3’ | 5’-cctcccctaccacaactctttgcc-3’ |
| | Coilin | 5’-gcagagacacaggttggc-3’ | 5’-gcgacaaatccaactacctgttaactc-3’ |
